# Synthetic standards combined with error and bias correction improves the accuracy and quantitative resolution of antibody repertoire sequencing in human naïve and memory B cells

**DOI:** 10.1101/284810

**Authors:** Simon Friedensohn, John M. Lindner, Vanessa Cornacchione, Mariavittoria Iazeolla, Enkelejda Miho, Andreas Zingg, Simon Meng, Elisabetta Traggiai, Sai T. Reddy

## Abstract

High-throughput sequencing of immunoglobulin repertoires (Ig-seq) is a powerful method for quantitatively interrogating B cell receptor sequence diversity. When applied to human repertoires, Ig-seq provides insight into fundamental immunological questions, and can be implemented in diagnostic and drug discovery projects. However, a major challenge in Ig-seq is ensuring accuracy, as library preparation protocols and sequencing platforms can introduce substantial errors and bias that compromise immunological interpretation. Here, we have established an approach for performing highly accurate human Ig-seq by combining synthetic standards with a comprehensive error and bias correction pipeline. First, we designed a set of 85 synthetic antibody heavy chain standards (*in vitro* transcribed RNA) to assess correction workflow fidelity. Next, we adapted a library preparation protocol that incorporates unique molecular identifiers (UIDs) for error and bias correction which, when applied to the synthetic standards, resulted in highly accurate data. Finally, we performed Ig-seq on purified human circulating B cell subsets (naïve and memory), combined with a cellular replicate sampling strategy. This strategy enabled robust and reliable estimation of key repertoire features such as clonotype diversity, germline segment and isotype subclass usage, and somatic hypermutation (SHM). We anticipate that our standards and error and bias correction pipeline will become a valuable tool for researchers to validate and improve accuracy in human Ig-seq studies, thus leading to potentially new insights and applications in human antibody repertoire profiling.

## INTRODUCTION

Adaptive immune responses are governed by cooperative interactions between B and T lymphocytes upon antigen recognition. A hallmark of these cells is the somatic generation of clonally unique antigen receptors during primary lymphocyte differentiation. In particular, B cell antigen receptors (BCRs, and their analogous secreted form, antibodies) result from rearrangement of the germline-encoded variable (V), diversity (D, heavy chain only), and joining (J) gene segments. V(D)J recombination in B cells creates a highly complex receptor population (generally interchangeably referred to as BCR, antibody, or immunoglobulin (Ig) repertoires), which matures upon antigen experience to produce the more targeted, high-affinity memory BCR network. In-depth and accurate characterization of these repertoires provides valuable insight into the generation and maintenance of immunocompetency, which can be used to monitor changes in immune status, and to identify potentially reactive clones for therapeutic or other uses. Due to rapid technological advances, high-throughput sequencing of Ig genes (Ig-seq) has become a major approach to catalog the diversity of antibody repertoires (1-3). Ig-seq applied to human B cells has potential in a variety of applications (4), particularly in antibody drug discovery (5-7), diagnostic profiling for vaccines (8, 9), and biomarker-based disease detection (10, 11). Additionally, Ig-seq is enabling a more comprehensive understanding of basic human immunobiology, such as B cell clonal distribution across physiological compartments in health and disease (12, 13).

A major challenge in Ig-seq is the requirement of accurate and high-quality datasets. Several current library preparation protocols are based on target enrichment from genomic DNA or mRNA (14). For example, the conversion of mRNA (more commonly used due to transcript abundance and isotype splicing) into antibody sequencing libraries relies on a number of molecular reagents and amplification steps (e.g. reverse transcriptase, multiplex primer sets, PCR), which potentially introduce errors and bias. Due to the highly polymorphic nature of repertoires especially from affinity-matured memory B cells and plasma cells, it becomes essential to determine if such technical noise occurs at non-negligible rates, as this could alter quantitation of critical repertoire features such as clonal frequencies, germline gene usage, and somatic hypermutation (SHM) (14, 15). One way to address this is through the use of synthetic control standards, for which the sequence and abundance is known prior to sequencing, thus providing a means to assess quality and accuracy (16). Several examples of standards have been presented for Ig-seq; Shugay et al. sequenced libraries prepared from a small polyclonal pool of B and T lymphocyte cell lines, and observed nearly 5% erroneous reads, resulting in approximately 100 false-positive variants per clone (17). Recently, Khan et al. developed a set of synthetic RNA (*in vitro* transcribed) spike-in standards based on mouse antibody sequences, which were used to show that a substantial amount of errors and bias are introduced during multiplex-PCR library preparation and sequencing (18).

Various experimental and computational workflows exist to mitigate the effects of errors and bias in Ig-seq. One of the most advanced and powerful strategies is to prepare libraries with the incorporation of random and unique molecular identifiers (UIDs, also commonly referred to as UMIs or molecular barcodes). Following sequencing, error correction can be performed by clustering and consensus building of reads that share the same UID; reads sharing the same UID are assumed to be derived from the same original mRNA/cDNA molecule (19). Furthermore, bias correction for cDNA abundance can be performed by counting the number of UIDs (instead of total reads) (20, 21). Several iterations of UID-tagging have been developed for Ig-seq, such as UID labeling during first- and second-strand cDNA synthesis (22), UID addition during RT template switching (23), and so-called “tagmentation” of UID-labelled amplicons (24). Recently, we developed an innovative strategy to add UIDs both during first-strand cDNA synthesis as well as multiplex-PCR amplification; this protocol, known as molecular amplification fingerprinting (MAF), results in comprehensive error and bias correction of mouse antibody repertoires (18).

Here, we describe an experimental-computational approach to generate highly accurate human Ig-seq data. We first designed a comprehensive set of synthetic standards based on human antibody heavy chain variable (IGHV) sequences: a total of 85 *in vitro* transcribed RNA standards, each with a unique complementarity determining region 3 (CDR3) sequence and covering nearly the entire set of productive human Ig heavy chain (IgH) germline (IGHV) gene segments. We used these synthetic standards to quantify the impact of errors and bias introduced during multiplex-PCR library preparation, and the robustness with which our previously developed method for UID addition by MAF could correct these artifact sequences. Finally, we implemented MAF-based error and bias correction on human B cell subsets (naïve and memory), which enabled us to make accurate clonal diversity estimates and quantify divergent repertoire features across B cell compartments.

## RESULTS

### Design of a comprehensive set of human synthetic standards

Our previously established set of murine synthetic antibody standards contained 16 unique clones (CDR3s) covering 7 IGHV gene segments (out of more than 140 annotated murine IGHV gene segments) (18). For our human standards, we developed a more comprehensive set consisting of 85 clones encompassing nearly the entire germline IGHV repertoire. The most commonly used repository for human germline segments is the International ImMunoGenetTics Database (IMGT), which has annotated 61 IGHV alleles as functional or having an open reading frame (25). After filtering out paralogs and selecting only gene segments that have been found in productive rearrangements (2), we chose 48 IGHV gene segments as the basis for our standards (**Table S1**). Each standard contained the following elements (5’ to 3’): (i) a conserved non-coding region, (ii) ATG start codon and a leader peptide sequence spliced to its respective IGHV gene segment, (iii) a synthetic CDR3 sequence, (iv) a germline IgH J (IGHJ) gene segment, (v) a non-coding synthetic sequence identifier (for the separation of standards from biological sequences), (vi) a partial segment of the constant region from isotypes IgM, IgG, and IgA (**Figure 1A**). This design allows amplification of our synthetic controls with a variety of PCR primer sets. Notably, for control singleplex-PCR experiments, all standards can be amplified by a single forward primer (targeting the conserved 5’ non-coding region) and a single reverse primer (targeting one of the isotype constant regions). Since IGHV gene segment usage has been reported to be non-uniform (2, 26), we selected the most abundant segments for use in multiple standards (**Figure 1B, Table S1**).

**Figure 1.**
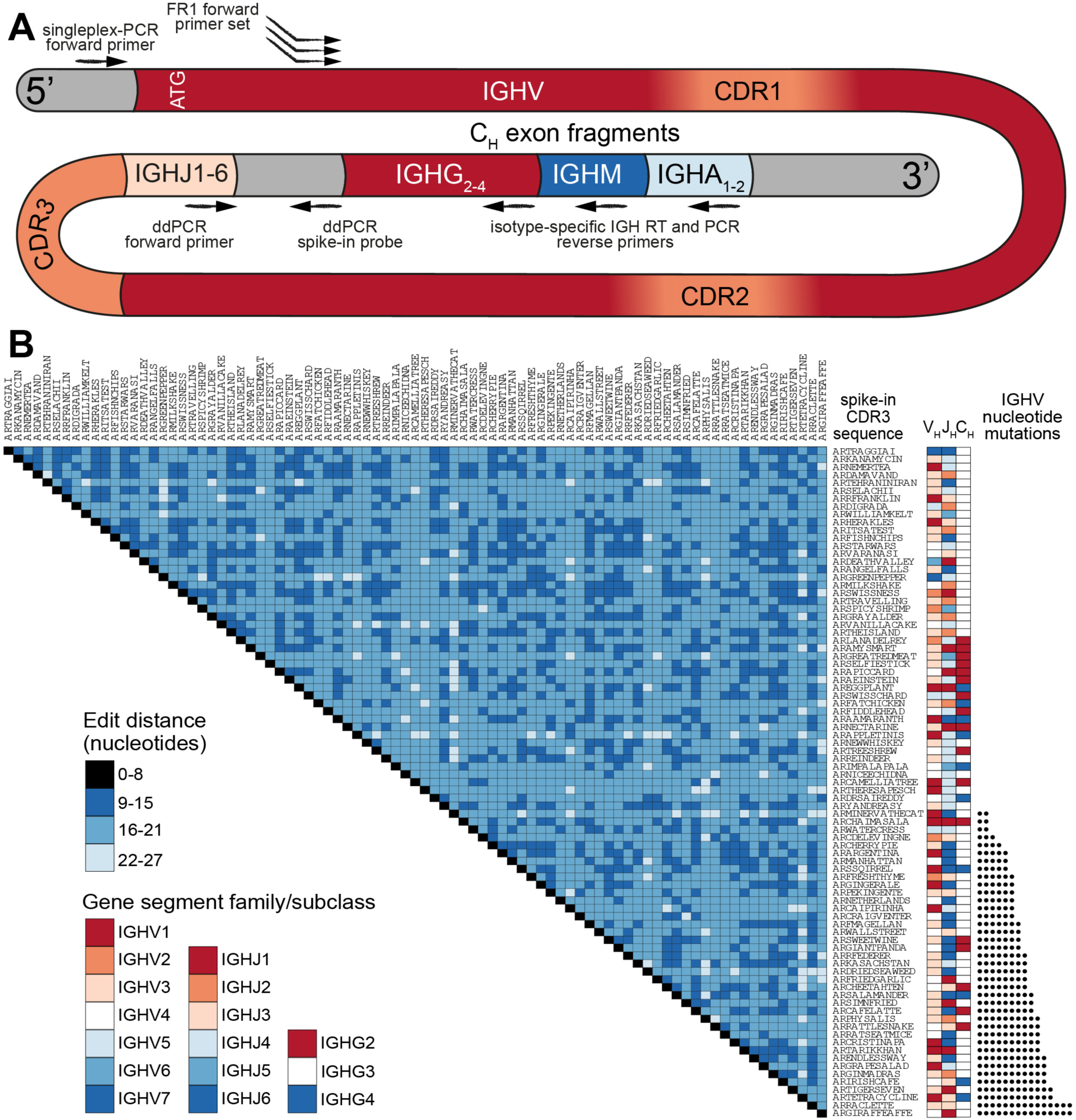
A comprehensive set of human synthetic spike-in standards for Ig-seq. **(A)** Schematic showing the prototypical spike-in with the following regions (5’ to 3’): a conserved non-coding region, ATG start codon, IGHV region with FR1 specific for multiplex-PCR, IGHJ regions, non-coding synthetic sequence identifier (specific for ddPCR probes), and downstream heavy chain constant domain sequences (IGHG, IGHM, IGHA) containing primer binding sites used for cDNA synthesis. Each spike-in contains a complete VDJ open reading frame, including nucleotides upstream of the ATG start codon, and downstream constant domain sequences containing primer binding sites used for sample cDNA synthesis. **(B)** Pairwise comparisons based on a.a. Levenshtein edit distance of all 85 standard CDR3 sequences. The germline IGHV and IGHJ segment family usage and IgG subclasses are denoted. 39 spike-ins contain rationally designed nt SHM (black circles) across the IGHV regions.

All standards carry a unique CDR3 sequence, which visually aids the analysis of sequencing results. Furthermore, all clones were designed to be resilient against sequencing and PCR errors: at least 9 specific nucleotide (nt) deletions, insertions, and/or mutations are needed in order to turn one CDR3 nt sequence into another (**Figure 1B**). For our experiments, synthetic standard genes were *in vitro* transcribed to RNA and subsequently reverse transcribed to cDNA. We measured individual cDNA molecules by digital droplet PCR (ddPCR) and capillary electrophoresis. Standards were then pooled in a non-uniform concentration distribution and maintained as a master stock (**Table S1**).

### Human Ig-seq library preparation using the MAF protocol

We adapted our previously described library preparation protocol for murine antibody repertoires to be compatible for human Ig-seq (18). This protocol is based on targeted amplification via RT of RNA to first-strand cDNA, followed by two PCR amplification steps (27, 28), the first of which uses a forward multiplex primer set targeting the IGHV framework region 1 (FR1). Each step also incorporates fragments of Illumina sequencing adapters (IA), such that the final product of the workflow is already compatible with the Illumina sequencing platform (**Figure 2A**).

**Figure 2.**
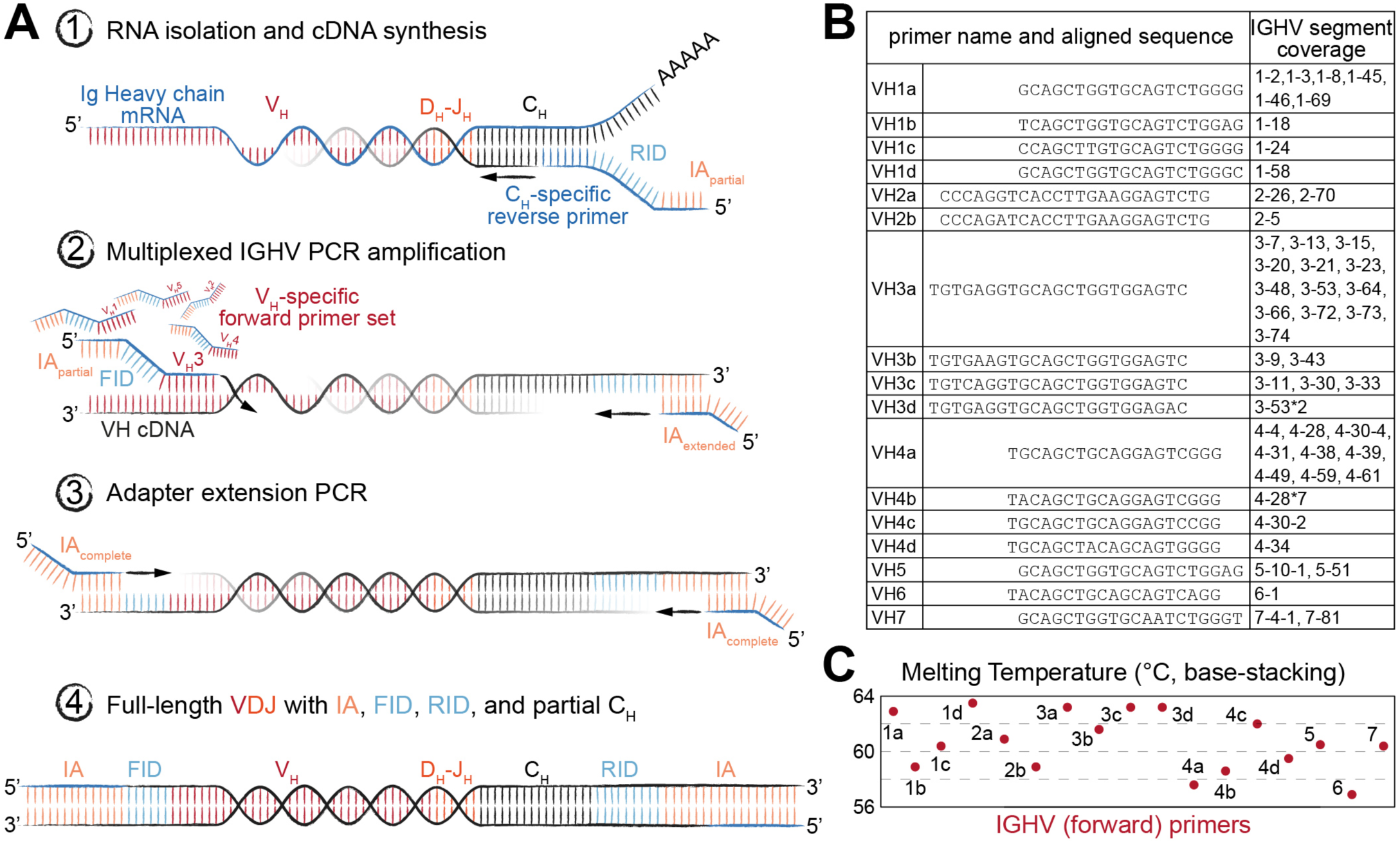
Library preparation of immunoglobulin (Ig) heavy chain genes for high-throughput sequencing (Ig-seq) using molecular amplification fingerprinting (MAF). **(A)** In step 1, reverse transcription (RT) is performed to generate first-strand cDNA with a gene-specific (IgM or IgG) primer which includes a unique reverse molecular identifier (RID) and partial Illumina adapter (IA) region. This results in single-molecule labeling of each cDNA with an RID. In step 2, several cycles of multiplex-PCR are performed using a forward primer set with gene-specific regions targeting heavy chain variable (V_H_) framework region 1 (FR1), with overhang regions comprised of a forward unique molecular identifier (FID) and partial IA. In step 3, singleplex-PCR is used to extend the partial IAs. The result (Step 4) is the generation of antibody amplicons with FID, RIDs, and full IA ready for Ig-seq and subsequent MAF-based error and bias correction. **(B)** List of oligonucleotides sequences annealing to the V_H_ FR1 used in multiplex-PCR (Step 2) of the MAF library preparation protocol. The nearest germline IGHV segment(s) likely to be amplified by the respective primer are listed in the rightmost column. **(C)** The estimated melting temperature distribution of the V_H_ FR1 forward primer set.

Importantly, our library preparation protocol used primers incorporating random-nucleotide UIDs, thus enabling MAF-based error and bias correction. A reverse-UID (RID) with theoretical diversity up to 2×10^7^ unique sequences is present in the RT primer (between the Ig constant region-specific and partial IA regions), and a forward-UID (FID with additional diversity of approximately 7x10^5^ unique sequences) is present on the forward multiplex primer set (between the FR1-specific and partial IA regions) used in the first PCR reaction. Such high diversity among RIDs is necessary in order to prevent tagging of different cDNA molecules with the same barcode. Our multiplex forward primer set was designed to target all IMGT-annotated IGHV gene segments (**Figure 2B**). To compromise between maintaining similar amplicon length across gene segment families and creating thermodynamically equivalent oligonucleotides, we placed the primers at or near the beginning of each FR1, resulting in a melting temperature range of 57°C to 63°C (**Figure 2C**). This range and the accompanying (unavoidable) variability in GC content have been shown to potentially cause differences in amplification efficiencies, which in turn leads to a biased representation of segment usage frequencies in Ig-seq data. Our workflow aimed to solve this problem in two ways: first, since the RID labels cDNA at the single molecule level, we are able to resolve the number of molecules by counting the number of RIDs instead of raw reads. Second, by using the FIDs on our forward primers, we are able to further normalize our molecular count, since Ig genes preferentially amplified by our primer set should show a higher ratio of FIDs to RIDs. Additionally, the RIDs can be used to correct for errors introduced during PCR and the sequencing process itself by grouping sequencing reads based on their RID, then correcting diverging nt positions by generating a consensus sequence (majority voting scheme). This is especially useful in Ig-seq when attempting to distinguish true SHM variants from erroneous sequences.

### Combining standards with MAF to correct errors and bias in Ig-seq

To evaluate the extent of errors and bias present in human Ig-seq data, standards were mixed (spiked-in) with cDNA prepared from circulating purified human B cells. In total, we sequenced 28 independently prepared libraries and annotated them with a custom aligner (18, 29). Prior to alignment, reads were either kept as uncorrected (raw) reads or were corrected using our MAF pipeline that takes into account RID and FID information. In this way, we could directly compare the number of erroneous sequences produced in uncorrected vs. MAF-corrected datasets. Clonal assignment of uncorrected reads produced many erroneous CDR3 amino acid (a.a.) variants (sequences with at least 1 a.a. difference from the nearest standard control sequence); for example, in a dataset with 100,000 aligned reads, there was a median value of 23 errors per clone (**Figure 3A**). The number of erroneous variants produced showed a clear correlation with the individual abundance of each clone within the master stock (***r*** = 0.89). When taking the entire VDJ nt sequence into account, an even greater number of erroneous variants were observed (≥1 nt difference from the standard sequence) (**Figure 3B**). We observed a median value of 118 erroneous nt variants per standard (per 100,000 aligned sequences). Again, the number of erroneous variants exhibited a clear correlation with clone abundance (***r*** = 0.90). However, we did not observe any significant trend linking IGHV family to the error rate (F-Test on full and reduced linear model, p = 0.083).

**Figure 3.**
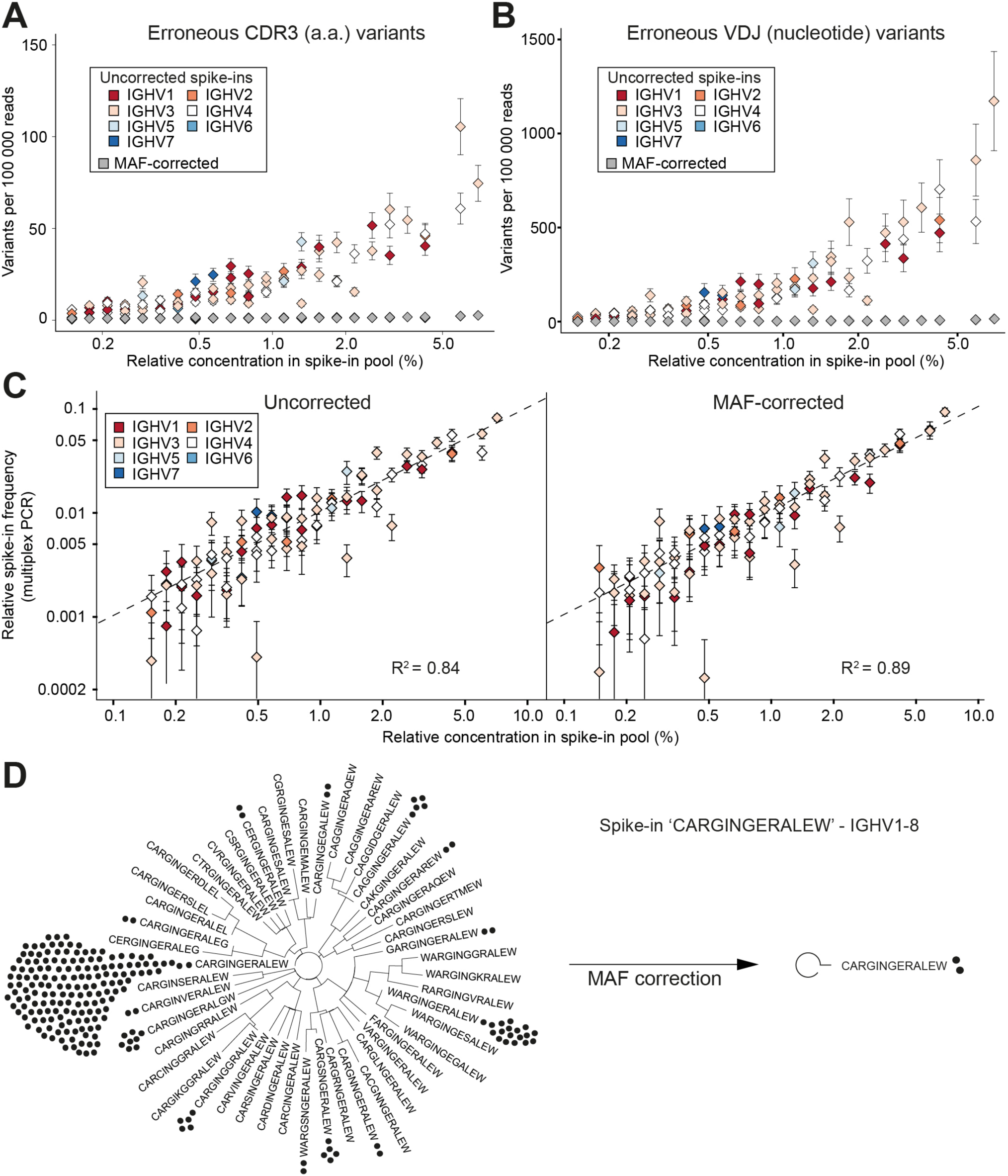
Synthetic standards used to validate performance of error and bias correction of Ig-seq data by MAF. **(A)** The number of erroneous CDR3 sequences (at least one a.a. difference from the correct CDR3) per 100,000 reads is plotted against the relative concentration of each standard (from a master stock). Color-coded diamonds correspond to germline IGHV segment family of the respective standard and show the number of erroneous variants in uncorrected (raw) data; gray diamonds indicate the number of variants remaining after MAF error correction. **(B)** The number of erroneous VDJ variants derived from each standard was calculated by finding all variants that carried the correct CDR3 a.a. sequence, but differed by at least one nt across the entire VDJ region. Colored diamonds represent uncorrected data; gray diamonds indicate variants remaining after MAF error correction. **(C)** Sequencing bias introduced by multiplex-PCR using the FR1 primer set was assessed by plotting the measured frequencies of each standard against its relative concentration (from a master stock). Dashed line represents a bias-free ideal scenario (R^2^ = 1). The left and right plots show observed frequencies before and after MAF bias correction, respectively. **(D)** Phylogenetic trees visualizing the CDR3 a.a. variants present for a selected standard with the CDR3 a.a. sequence *CARGINGERALEW* and IGHV1-8 and IGHJ1 segment usage. Prior to error correction, 39 erroneous CDR3 a.a. variants (branches) and 218 VDJ nt variants (black circles) were observed. Following MAF error correction, only the original correct CDR3 a.a. and two VDJ nt variants remain. The Ig-seq data sets used in A-C consisted of ∼300,000 preprocessed full-length antibody reads from each of the synthetic spike-in only samples. IgG1_D1 dataset was used for panel **(D)** (see **Table S3**).

After performing error correction with our MAF pipeline, there was a dramatic reduction of CDR3 and VDJ errors: we observed a median value of 0 and 1 error per clone, respectively. Across all datasets, we remove an average of 94.2% CDR3 a.a. and 97.4% VDJ nt erroneous variants (**Figure 3A-B**). For example, prior to error correction the standard “CARGINGERALEW” (from dataset Donor 1, IgG aliquot 1, see **Table S3**) displayed 39 additional CDR3 a.a. variants and 217 additional VDJ nt variants (**Figure 3D**). After error correction, we retain only the correct CDR3 a.a. sequence and only one additional (erroneous) nt variant. With our current filtering criteria (see Methods), we observed 16 instances in which we removed a standard control sequence that was present in the raw data. However, in the vast majority of cases, we kept the standard when it was observed in the raw data (2,321 instances). In only 43 instances, a standard sequence was either too low in abundance or too frequently mutated to be annotated in either the raw or error-corrected datasets.

In order to assess potential biases introduced by library preparation we prepared control libraries containing only the pool of synthetic standards (from the master stock). The libraries were generated in the same manner as the described MAF protocol, with the exception that in the first PCR step, instead of using a multiplex forward primer set, a single forward primer targeting the conserved 5’ non-coding region (singleplex-PCR) was used. Ig-seq on these samples allowed us to establish a baseline for pipetting accuracy by comparing the obtained standard frequencies from the singleplex-PCR against the expected frequencies based on our pooling scheme: this yielded an R^2^ of 0.88 and average mean squared error (MSE) of 0.29 ± 0.02% (**Figure S1A**). These values indicate that only small systematic deviations occurred, most likely due to minor pipetting error. Next, we compared standard frequencies (expected relative concentration) with frequencies generated in our previous multiplex-PCR libraries, both with and without MAF correction (**Figure 3C**). On uncorrected data, the multiplex-PCR libraries achieved an R^2^ of 0.84 with an average MSE of 0.34 ± 0.06%, which is significantly worse than the value obtained by singleplex-PCR (Student’s t-test p = 0.008). After error and bias correction on these same datasets, the correlation improved to an R^2^ of 0.89 and an average MSE of 0.28 ± 0.08%. While MAF-corrected MSE values show no significant difference to the singleplex-PCR libraries (Student’s t-test p = 0.49), they do highlight a significant improvement over the uncorrected data (paired Student’s t-test, p = 0.0007).

### Impact of MAF error correction on human B cell repertoires

Next, we analyzed the impact of MAF on the BCR repertoires of B cells isolated from the peripheral blood of three healthy donors. We used flow cytometry sorting and a gating strategy to select for CD27^-^ IgM^+^ (naïve) and CD27^+^ IgG^+^ (memory) B cell populations (**Figure 4A**). Across all donors, the fraction of CD19^+^ peripheral blood B cells was 16-29% for CD27^-^ IgM^+^ and 5-9% for CD27^+^ IgG^+^. Importantly, each donor population was split into 4-5 separate aliquots containing 200,000 cells each (cellular replicates) prior to cell lysis. Total RNA was extracted and RT for cDNA synthesis was performed independently in order to prevent the mixing of transcripts across cellular replicates. The cDNA of synthetic standards (from the master stock) was then mixed with the B cell cDNA, and corresponding molecular quantities were measured by ddPCR (**Figure 4B, Table S3**). All cDNA libraries were then processed into libraries using the MAF protocol (**Figure 2A**) and subjected to Ig-seq.

**Figure 4.**
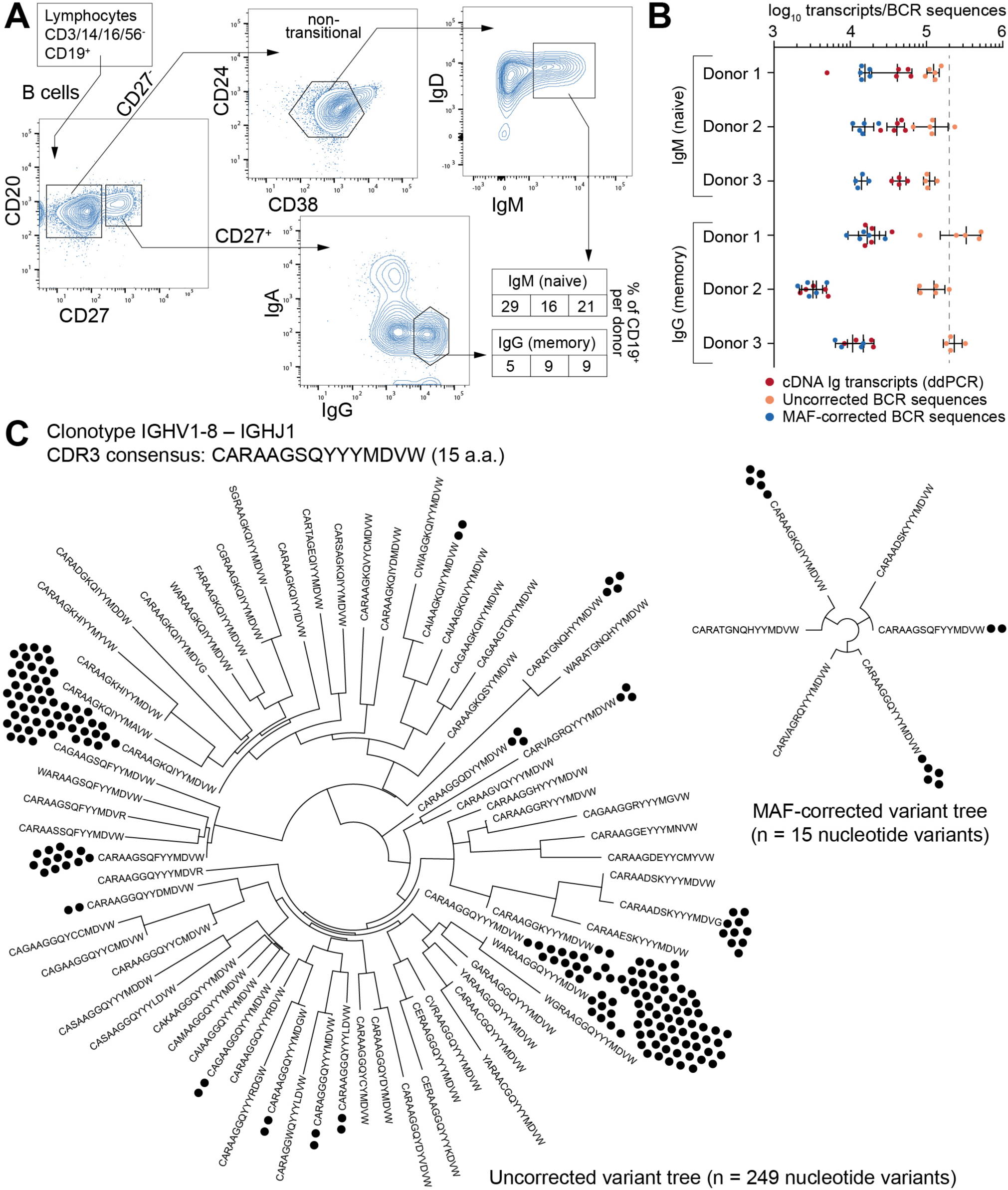
Ig-seq analysis of human naïve (CD27^-^IgM^+^) and memory (CD27^+^IgG^+^) B cells. **(A)** The flow cytometric workflow for isolating CD27^-^IgM^+^ and CD27^+^IgG^+^ B cells from peripheral blood. Boxed-in values indicate the frequency of each sorted subset as a percentage of the total B cell (CD19^+^) population from each of three donors (1-3 from left to right). **(B)** Experimental and Ig-seq based quantitation of antibody diversity; points represent cDNA molecule counts (using ddPCR) or unique reads (before and after MAF error correction) from cellular replicates (with mean and standard deviation shown) isolated from each donor and B cell subset. Unique read counts were based on the VDJ nt seqeunce. Dashed line represents the number of B cells isolated per cellular replicate (2 x 10^5^ cells). **(C)** Phylogenetic trees illustrating CDR3 a.a. and nt variants present for the selected clonotype with the consensus CDR3 sequence *CARAAGSQYYYMDVW* and IGHV1-8 to IGHJ1 recombination. Prior to error correction, 70 erroneous CDR3 a.a. variants and 249 VDJ nt variants (black circles) were observed. Following MAF error correction, only 6 CDR3 a.a. and 15 VDJ nt variants remain. The Ig-seq data sets used in **(B)** are described in **Table S3**; IgG1_D1 was used for the tree in panel **(C)**.

A simple global analysis of Ig-seq data revealed that diversity measurements were dramatically exaggerated, as the number of unique antibody sequence variants obtained from the raw, uncorrected data often exceeded both the number of cells and the number of total cDNA transcripts in a given aliquot. Following error correction by MAF, the variant count returned to ranges that are physically possible, thereby highlighting the importance of proper error correction and quality control when globally determining repertoire diversity (**Figure 4B**). We further examined the influence of erroneous variants on CDR3 clonotype analysis. In order to identify clonotypes, we used hierarchical clustering (30) based on sequences sharing the following features: identical IGHV and IGHJ gene segment usage, identical CDR3 length, and a CDR3 a.a. similarity of at least 80% to one other sequence in the given clonotype. When performing such an analysis on uncorrected data, clonotypes contained an artificially high number of distinct clones (**Figure 4C**, left tree). Here, the IgG-derived clonotype with the consensus CDR3 of ‘CARAAGSQYYMDVW’ (from the same sample and IGHV gene segment used by the standard shown in **Figure 3D**) contains 249 unique nt variants and 70 unique a.a. sequences. After MAF-based error correction, only 15 nt variants and 6 distinct CDR3 a.a. sequences remained. It is worthy to note that although both the standard sequence (**Figure 3D**) and the biological clonotype (**Figure 4C**) had a large number of CDR3 a.a. variants in uncorrected data, after error correction only the biological clonotype retained multiple a.a. variants, suggesting these may be true variants generated *in vivo* by SHM. This general trend of each IgG memory B cell-derived clonotype to contain more variants relative to antigen inexperienced IgM-expressing B cells was clear across our biological data sets (**Figure 5A**).

**Figure 5.**
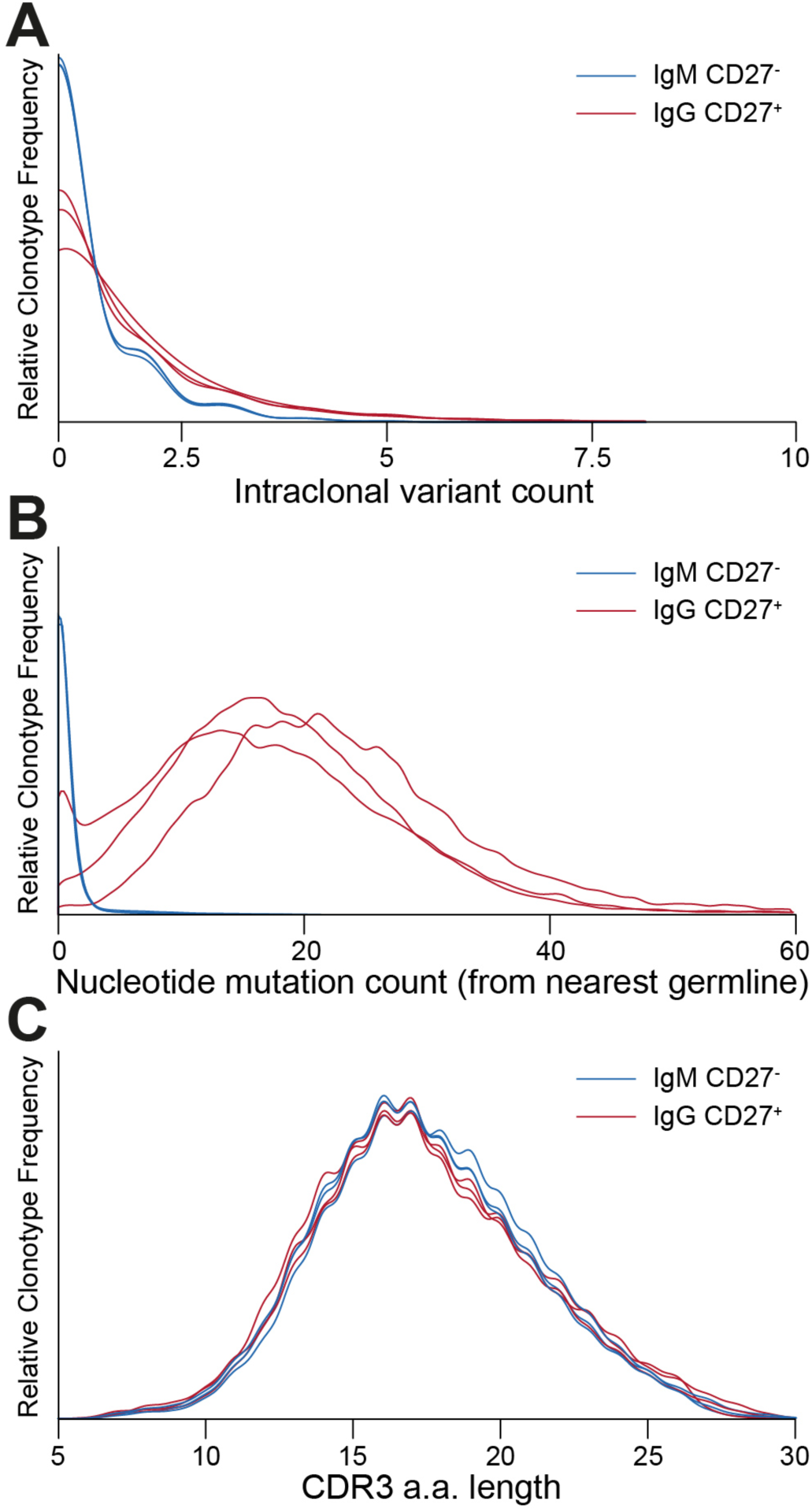
Ig-seq analysis of molecular features highlight global differences between naïve (CD27^-^ IgM^+^) and memory (CD27^+^IgG^+^) B cells. **(A)** Clonotype size (calculated as the total number of variants within a clonotype) for naïve IgM (blue lines) and memory IgG (red lines) B cells. Each pair of lines represents a single donor. **(B)** Graph showing the distribution of average SHM frequencies for VDJ nt variants per clonotype. The CD27^-^IgM^+^ B cell repertoires (blue lines) have a median SHM value of zero, whereas only a small fraction of clonotypes (approximately 8%) contain an average of one or more mutations. CD27^+^IgG^+^ repertoires (red lines) have a median of 20 to 24 SHM per nt variant within each clonotype. **(C)** CDR3 a.a. length distribution across clonotypes from naïve (blue lines) and memory (red lines) B cell subsets.

### Clonal diversity measurements of human B cell repertoires after error and bias correction

After establishing the value of performing MAF error correction on biological repertoires, we next focused on determining the clonotype diversity present in each B cell sample. First, we determined the overlap of clonotypes present in each cellular replicate (**Figure 6A**). Notably, we observed an overlap of several clonotypes in the CD27^-^IgM^+^ subset; this overlap was unexpected given that this subset should be highly enriched for naïve B cells, which by definition are not antigen experienced or clonally expanded, and should therefore be mostly unique (not present in multiple replicates). For each donor in the CD27^-^IgM^+^ subset, 1-2% of all clonotypes were present in at least one other cellular replicate. In the CD27^+^IgG^+^ subset, clonotypes shared between at least two cellular replicates were nearly tenfold more frequent (12-15%), which was expected given that this population is comprised of antigen experienced, clonally expanded memory B cells. Another observation discordant with the expected naïve B cell properties of the CD27^-^IgM^+^ subset was that overlapping clonotypes (in donors 1 and 3) were significantly more likely to have acquired mutations (**Figure 6B**), which are not a typical feature of naïve B cells. In comparison, over 90% of all CD27^+^IgG^+^ (overlapping and non-overlapping) clonotypes possessed at least one SHM, an expected observation in a memory subset.

**Figure 6.**
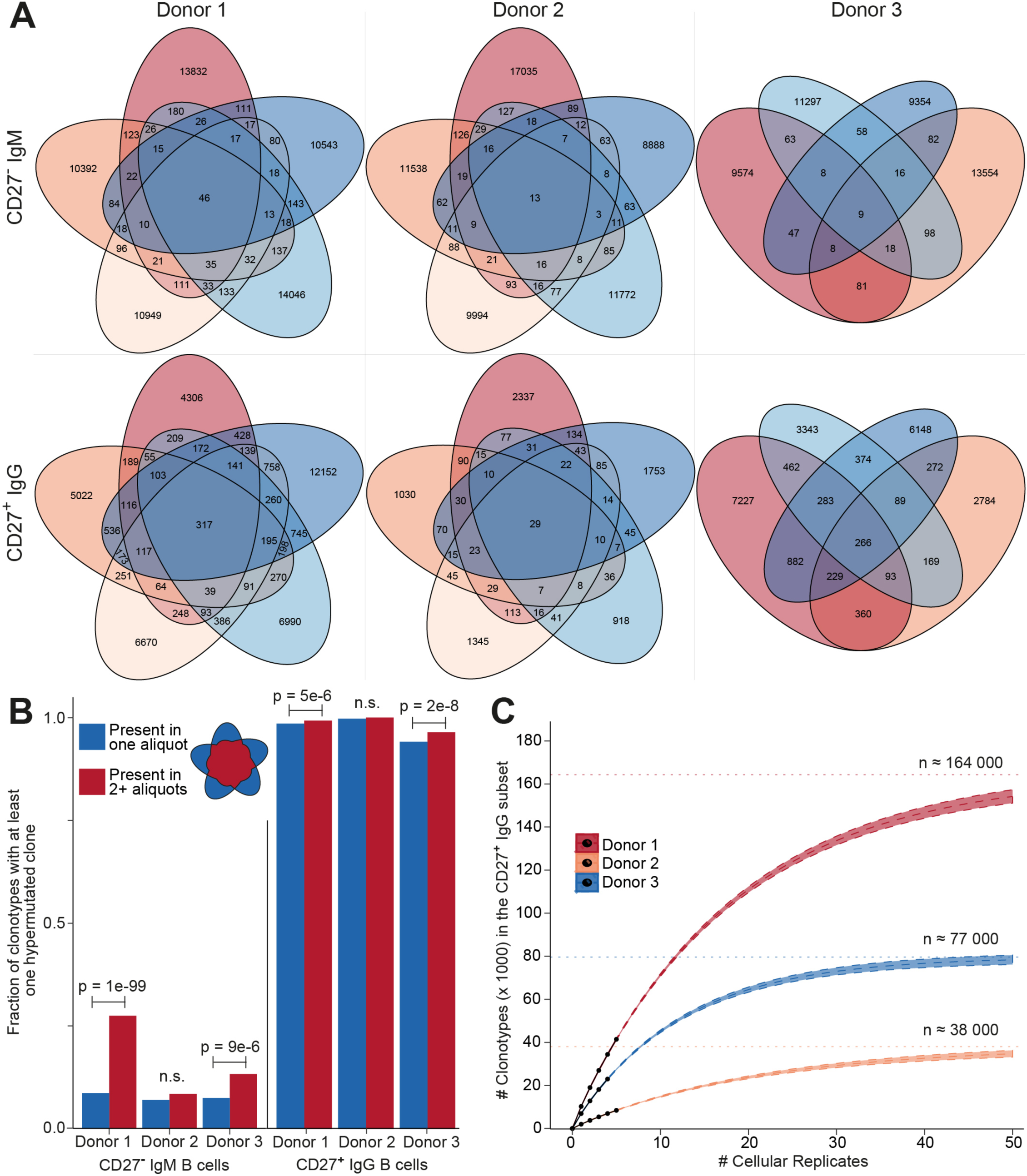
Clonotype diversity analysis across cellular replicates of naïve (CD27^-^IgM^+^) and memory (CD27^+^IgG^+^) B cells. **(A)** Venn diagrams show the presence of clonotypes (80% CDR3 a.a. similarity to least one clone in the cluster, same CDR3 a.a. length, same IGHV and IGHJ gene segment usage) across cellular replicates (2 x 10^5^ cells each) from each donor B cell subset. **(B)** Bar graph showing the fraction of clonotypes containing at least one variant with at least one nt SHM. Blue and red bars indicate clonotypes identified either in only one aliquot (unique) or in several aliquots (shared), respectively. The p-values represent significance using Fisher’s exact test. **(C)** Species accumulation curves for CD27^+^IgG^+^ B cells: the number of newly discovered clonotypes from each additional cellular replicate (black circles) is plotted. Extrapolating the observed overlap provides an estimate for the total number of distinct clonotypes (Chao2 estimator: D1 = 164,268 ± 2,365; D2 = 38,034 ± 1,302; D3 = 76,904 ± 1,409) and the approximate amount of cellular replicates needed to discover all clonotypes present in the peripheral blood CD27^+^ IgG^+^ population.

The high amount of overlap within the CD27^+^IgG^+^ B cell replicates of each donor allowed us to use established population diversity estimation techniques to calculate clonal diversity (31). Rarefaction curves were generated and estimates were extrapolated as a function of real and predicted cellular replicates (**Figure 6C**). The asymptote was determined by the standard form of the Chao2 estimator and yielded the following values for clonotype numbers: donor 1 = 164,268 ± 2,365, donor 2 = 38,034 ± 1,302, and donor 3 = 76,904 ± 1,409. Since the 95% confidence intervals for the three donors did not overlap, we could also infer that the size of each donor’s repertoire at the collection time point was significantly different. This analysis indicates that we would need to sample at least tenfold more cellular replicates in order to observe > 90% of all clonotypes; however the first five samples analyzed here were sufficient to observe > 25% of the clonotypic memory repertoire. We also generated rarefaction and extrapolation curves rescaled to the RID count (**Figures S2A and S2B**). In the case of the CD27^-^IgM^+^ repertoire data, while asymptotic curves could be generated, a diversity estimation is impractical. This is because plotting the observed numbers of newly discovered clonotypes for each additional RID and donor shows that the number of newly discovered clonotypes in the CD27^-^IgM^+^ dataset continues to grow over the observed range, whereas the number of new clonotypes starts to converge at approximately 20,000 RIDs for CD27^+^IgG^+^ repertoires (**Figure S2C**).

### Divergent features of CD27^-^IgM^+^ and CD27^+^IgG^+^ repertoires

After pooling all clonotypes (expanded and unique to a single cellular replicate) for each donor, we globally characterized sequences of the naïve CD27^-^IgM^+^ and memory CD27^+^IgG^+^ subsets. First, we determined the SHM count (nt) of each clone with respect to its nearest germline IGHV and IGHJ gene segment sequence. The median values of SHM for the CD27^-^IgM^+^ repertoires were zero, which was to be expected for a naïve B cell subset. In contrast, the median values for CD27^+^IgG^+^ repertoires were 20-24 mutations per clone (**Figure 5B**). It is widely appreciated that human heavy chain CDR3 sequences are much longer than their murine counterparts, which we also observed here, with a slight (but consistent across donors) variation between naïve and memory B cells (**Figure 5C**). Interestingly, the IGHV gene segment family usage correlated with B cell subset. The CD27^+^IgG^+^ repertoires across all donors were relatively enriched for IGHV1 and IGHV3 gene segment family members, whereas the relative share of the IGHV4 gene segment family was larger in the CD27^-^IgM^+^ repertoires (**Figure 7A**). We validated these observations quantitatively using linear discriminant analysis (LDA) fitted on the centered log ratio (CLR)-transformed frequencies of each cellular replicate (**Figure S3**). The LDA classifier was fit on different splits of the data (based on two of the donors) and used to predict a holdout set (based on the remaining donor). This showed that the fitted classifier in each instance is highly predictive of the remaining aliquots and that prediction is robustly driven by the relative abundance of IGHV4 segment family usage in the CD27^-^IgM^+^ repertoires and the IGHV1, 2, and 3 families in the CD27^+^IgG^+^ repertoires (**Figure S3A-C**). Next, we utilized LDA to perform dimensionality reduction of all data points to into a single one-dimensional axis; again, the most important components were the relative abundance of IGHV3 and IGHV4 gene segment families (**Figure S3D**).

**Figure 7.**
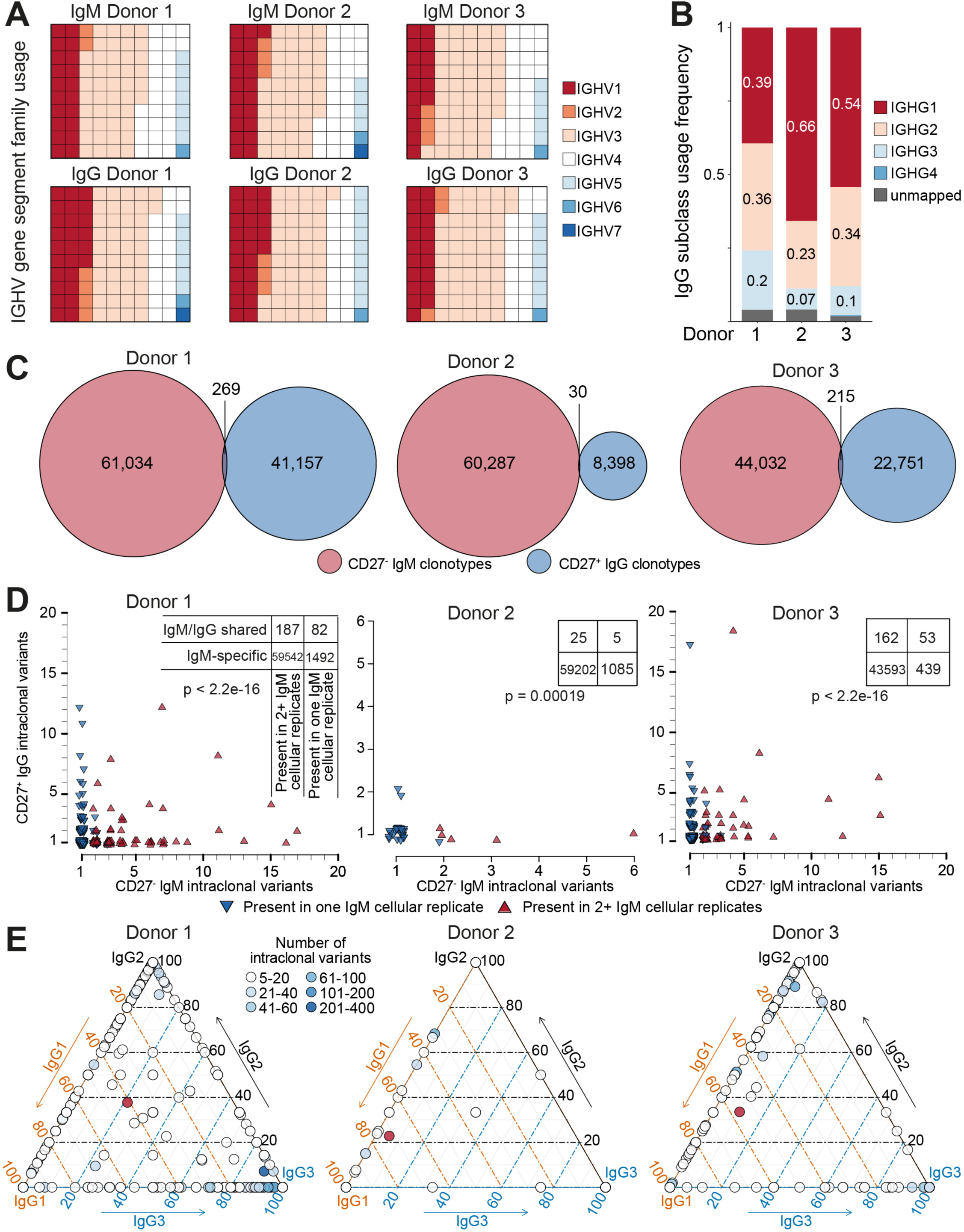
Genetic features of naïve and IgG memory BCR repertoires. **(A)** Block map shows IGHV gene segment usage sorted by family (color-coded) across donors and B cell subset; blocks are normalized to the total number of clonotypes within each group. **(B)** IgG subclass usage in CD27^+^IgG^+^ donor repertoires. IgG_4_ sequences (dark blue bars) were virtually absent among three donors; a small fraction of sequences could not be unambiguously mapped based on the sequencing read (gray bars). **(C)** Venn diagrams showing the overlap of clonotypes (80% CDR-H3 amino acid similarity to least one clone in the cluster, same CDR3 a.a. length, same IGHV and IGHJ gene segment usage) between the naïve (CD27^-^IgM^+^) and memory (CD27^+^IgG^+^) BCR heavy chain repertoire in each of three donors. **(D)** Each plot shows the clonal composition of each shared clonotype (from panel **(C)**) in terms of its IgG and IgM intraclonal variants. Red triangles indicate clonotypes found in multiple IgM cellular aliquots; blue triangles show clonotypes which could only be found in one IgM cellular aliquot. The total number of clonotypes found are depicted in the corresponding contingency table. Fisher’s exact test was used to quantitatively analyze enrichment of expanded IgM clonotypes in the shared IgM/IgG subset. **(E)** Ternary plots comprised of three axes representing the IgG1, IgG2, and IgG3 isotype subclasses. The relative subclass composition of intraclonal variants per IgG clonotype (each represented by a circle colored according to the number of variants belonging to that clonotype) is depicted by the position of the circle within the triagonal space. Red circles represent the average subclass composition for all IgG variants of each donor (cf. **Figure 6C**).

Next, we leveraged the ability of our reverse primer to distinguish among IgG subclasses (**Figure 7B**). The majority of sequences mapped either to IgG1 (40-66%), IgG2 (23%-36%), or IgG3 (23%-36%), whereas IgG4 sequences were extremely rare, observed solely in donor 3 (0.3%). Finally, we compared the CD27^-^IgM^+^ and CD27^+^IgG^+^ repertoires of each donor to determine the clonotype overlap of each B cell subset and isotype. Strikingly, the observed overlap was very small, with only 269 shared clonotypes for donor 1, 30 shared clonotypes for donor 2, and 215 clonotypes for donor 3 (**Figure 7C**). In each donor, these represented less than 0.5% of identified clonotypes. Closer examination revealed that clonotypes shared between the CD27^-^IgM^+^ and CD27^+^IgG^+^ subsets were also significantly enriched for intraclonal variants (SHM in CDR3) in one of the two populations (**Figure 7D and Figure S4**). Furthermore, we could see that clonotypes with multiple IgM variants were also shared specifically among IgM cellular replicates (**Figure 7D**, contingency tables). This intraclonal variant bias was not limited to heavy chain isotype: IgG clonotypes with ≥ 5 intraclonal variants also exhibited subclass composition skewing toward the IgG1/2 or IgG1/3 axis, but rarely at proportions similar to the overall IgG subclass distribution (**Figure 7E**).

## DISCUSSION

Ig-seq is becoming an essential tool for the quantitative analysis of antibody repertoire diversity and distribution. However, similar to other areas of high-throughput sequencing, Ig-seq also suffers from technical errors and bias; thus, standardized experimental and analytical methods that increase the validity of immunological interpretations must be developed. Here, we establish a comprehensive set of synthetic standards which, when combined with UID-labeling and MAF-based error and bias correction, results in highly accurate antibody repertoire data. By applying this approach to human B cell subsets, we gain unique insights into repertoire features such as clonal diversity, germline gene usage, SHM, and clonal history.

The synthetic standards developed here allowed quantitative interrogation of several accuracy-related features in Ig-seq. One major observation was that raw uncorrected data has a high number of erroneous variants, found both within the clonotype-defining CDR3 and across the entire VDJ region (**Figure 3A, B**). The number of false-positive variants correlated with the abundance of each standard; this is of particular concern because high frequency, clonally expanded B cells are often correlated with antigen specificity and thus important for biological interpretations (32). However, when applying our MAF-error correction protocol, we were able to remove nearly all erroneous CDR3 and VDJ variants (94-97%); this correction was robust even for high-frequency standards where the number of erroneous variants was especially high. Errors not removed by MAF could potentially be addressed with more stringent filtering criteria (e.g. read number cutoffs); however, this may come at the cost of reducing overall dataset size and removing legitimate intraclonal variants in biological samples.

Another aspect we quantified with our standard pool was the impact of multiplex primer sets, which have been shown to introduce substantial bias during library preparation (18, 33). By designing our standards with a 5’ conserved singleplex region (**Figure 1A**), we were able to directly compare Ig-seq data from libraries (on the same master stock) prepared by singleplex-PCR vs. multiplex-PCR. Our newly designed FR1-targeting multiplex primer set (**Figure 2B, C**) demonstrated a relatively strong correlation with singleplex-PCR data (R^2^ = 0.84) (**Figure 3C**). However, by performing multiple MAF error and bias correction steps, the correlation was improved by an additional 7% (**Figure 3D)**. The remaining variability does not appear to be restricted to a particular IGHV-gene family, indicating there is little systematic bias with respect to homologous sequences within the standard pool. MAF therefore represents an essential step in eliminating technical artifacts from human Ig-seq workflows, as it is able to generate data that closely mirrors that of the original sample. In future applications, these synthetic standards could be a critical asset for evaluating newly designed primer sets, library preparation protocols, or implementing new error and bias correction pipelines.

Having established a comprehensive set of synthetic standards and a validated error and bias correction pipeline, we were able to perform several analyses on human B cell repertoires with greater confidence in the accuracy and quantitative resolution of the Ig-seq data. A simple approach to estimating repertoire diversity is to calculate the number of unique antibody sequences as a fraction of total transcript (cDNA) input. However, performing this analysis on our samples suggests that the CD27^+^IgG^+^ memory B cell compartment is significantly more diverse than the naïve CD27^-^IgM^+^ B cells (82% vs. 35% unique nt variants, respectively, averaged across donors and cellular aliquots, Student’s t-test p < 10^-4^). This is potentially due to sample size variability with respect to the number of transcripts and our ability to oversample smaller libraries. Critically, when using bulk-sorted cells with UID labeling, it is not possible to discriminate between transcript copies that are identical because they came from the same lysed cell, and those which are identical because they represent two distinct, clonally related B cells. Thus, by biological subsampling through cellular replicates, we ensured that clonotypes observed in multiple samples must come from distinct, clonally related B cells, thereby providing an effective solution for estimating clonal diversity.

Applying computational approaches from ecology (31) to our biological subsampling strategy, we attempted to estimate the number of unique clonotypes in a given antibody repertoire. The CD27^-^ IgM^+^ B-cell subset did not show substantial clonotype overlap among cellular aliquots (**Figure 6A**). As it is commonly assumed (and typically the case as shown in **Figure 5A**) that each newly generated B cell is a unique clone, the size of the naïve repertoire in the human peripheral blood would be equal to the total number of naïve B cells, in the range of 10 to 30 million. While it is improbable to sample this subset in its entirety, and its diversity is also too high to estimate based on the cell numbers obtained here, our observations are consistent with this model, since each additional cellular replicate produced overwhelmingly unique sequences. One donor did show an unexpected presence of overlapping sequences (shared clonotypes) across IgM cellular replicates (**Figure 6A**, donor 1); these clonotype sequences were significantly enriched for SHM (**Figure 6B**), suggesting the possible presence of an antigen-experienced B cell subset within CD27^-^IgM^+^ population, and highlighting the need for improved characterization of the heterogeneity within circulating human B cell subsets. In the CD27^+^IgG^+^ B cell subset, we observed substantially more overlap across cellular replicates, which was expected given that memory-enriched B cells would have experienced antigen and undergone clonal expansion. By extrapolating the numbers of additional uniquely observed clonotypes with each subsequent cellular aliquot, we were able to predict the clonotype size of the peripheral CD27^+^IgG^+^ B-cell repertoire to be on the order of 10^5^ (**Figure 6C**). Indeed, rough estimates of the number of antigen-specific clonotypes generated by a single immune response (≈100, a number in line with what has been described regarding serum antibody clonotypes (34)) and the number of structurally distinct pathogens against which an individual has mounted a response (≈1000, a generous estimate given work showing that worldwide, individuals have on average a serological history against less than 100 viral species (35)) suggest that a memory repertoire of this size could reasonably protect against latent infection and/or subsequent antigen encounter.

We observed a clear shift in IGHV segment family usage from the naïve to the memory BCR repertoire (**Figure 7A**). Consistently observing this reshaping in three independent healthy donors, and comparing to our standard controls to exclude the possibility of biased amplification, we can conclude that it is a genuine phenomenon. Relatively more abundant IGHV1 and less abundant IGHV4 segment usage in IgG memory B cells has been previously observed in one three-donor cohort (1) but not in another which pooled sequences from both class-switched and IgM-memory cells (36), underscoring the importance of experimental design and accurate bias correction in antibody repertoire analysis.

Our Ig-seq workflow also allowed us to unambiguously assign IgG antibody sequences to their appropriate subclass, offering further insight into patterns of class-switch recombination present in memory-enriched B cells. While plasma cell-secreted IgG proteins in human serum are present at ratios of approximately 14:8:1:1 (for IgG1:2:3:4, respectively (37)), CD27^+^IgG^+^ B cells showed a distribution of approximately 5:3:1 for IgG1:2:3, with a nearly complete absence of IgG4 (**Figure 7B**). Cole et al. similarly observed a lack of IgG4 heavy chains in a single donor but described an enrichment of IgG2 relative to IgG1 and IgG3 (24). The abundance of IgG3^+^ B cells relative to its presence in the serum seen here indicates IgG3 may play a more important role in maintaining the reactive memory response compared to the protective memory response provided by serum IgG. Notably, the IgG3 locus is the most proximal, and thereby the most plastic of the human IgG subclasses; that is, an IgG3^+^ B cell still retains the capacity to class-switch to any of the remaining three IgG subclasses, whereas IgG1, IgG2, and IgG4 cannot return to any of the previous states. Similar to these findings, a flow cytometry-based investigation has also found healthy human donors to have low frequencies of IgG4-expressing circulating memory B cells (38).

With new daily production and relatively rapid turnover of naïve B cells, it was not unexpected to see little overlap of clonotypes between the intradonor CD27^-^IgM^+^ and CD27^+^IgG^+^ populations (**Figure 7C**). An interesting finding was that for clonotypes present in both B cell subsets, intraclonal variation was largely restricted to one of the two isotypes. Assuming that clonotype overlap among CD27^-^IgM^+^ cellular replicates represents the presence of antigen-experienced, clonally expanded B cells, this suggests that the antigen specificity of antibody variable domains may to some extent be influenced by the downstream constant regions, which has been observed functionally for small cohorts of human and murine IgG and IgA antibodies (39). Notably, we also observe similar clonal restriction within IgG clonotypes with respect to heavy chain subclass (**Figure 7E**). This may be driven by the type of antigen and the nature of the elicited immune response, or governed by physical constraints as we suggest for the differences between the IgM and IgG repertoires. A larger scale functional study, including IgM sequences, could provide crucial support for this model, which would shed new light on the role of Ig isotypes and subclasses on B cells in the post-antigen encounter setting.

## METHODS

### Preparation of spike-in master stocks

The spike-in standards were ordered from GeneArt (Invitrogen) in the form of plasmids. Each spike-in sequence contained a T7 promoter for *in vitro* transcription. Approximately 1.5 µg of each plasmid was digested with 10 U of EcoRV-HF (New England BioLabs) and purified with DNA-binding magnetic beads (SPRI select, Beckman Coulter). Approximately 1 µg of the digested plasmid was then used for *in vitro* transcription (MEGAscript T7 Transcription Kit, ThermoFisher Scientific). RNA was purified by lithium-chloride precipitation, eluted (TE with 1U/µl RiboLock) and aliquoted. The final concentration was then determined with the TapeStation (Agilent Technologies).

The spike-in RNA obtained this way was reverse transcribed with the Maxima Reverse Transcriptase kit (Thermofisher Scientific). 500 ng mRNA was mixed with 20 pmol of IgM reverse primer and 3 µl dNTP-mix (10 mM each) and was then filled up to 14.5 µl with water. The reaction-mix was incubated at 65 °C for 5 minutes. 4 µl of 5x RT-buffer, 0.5 µl (20 U) RiboLock and 1 µl Maxima reverse transcriptase (200 U) were then added. The resulting reaction mix was incubated for 35 minutes at 55 °C, followed by a termination step at 85 °C for 5 minutes. 2.5 µl of RNase A (Thermofisher Scientific) was added and the mix was again incubated at 60 °C for 30 minutes. The resulting cDNA was purified with SPRI Select magnetic beads and eluted in nuclease-free water. The concentration of each cDNA reaction was determined afterwards with the Fragment Analyzer and pooled according to Table S1. The exact concentration of the pooled spike-ins was determined by ddPCR with dilutions of the pool ranging from 10^-3^ to 10^-6^. The measured spike-in pool was afterwards diluted to a final storage concentration of 250,000 transcripts per µl.

### Transcript quantitation by ddPCR

Quantifiying cDNA and PCR products by ddPCR was conducted for all measurements in the following way: A dilution series with 3 or 4 points was prepared. Droplets were generated with BioRad’s droplet generator using 12.25 µl of ddPCR Supermix (BioRad) combined with 10 µl of the diluted sample, 25 pmol of the biological ddPCR probe (SF_21), 25 pmol of the spike-in specific ddPCR probe (TAK_499), 22.5 pmol of the forward (SF_63) and reverse ddPCR primer (TAK_522) and 55 µl of droplet generation oil (BioRad). Droplets were then transferred to a 96-well reaction plate, which was heat sealed with easy pierce foil (VWR International). Then a PCR reaction was performed using the following conditions: 95°C for 10 min; 45 cycles of 94 °C for 30s, 53 °C for 30s, 64 °C for 1 min; 98°C for 10 min; and holding at 4 °C. After the PCR step, every 96-well plate was read using BioRad’s droplet reader.

### B-cell isolation, sorting, and lysis

Peripheral blood leukocyte-enriched fractions (‘buffy coats’) were received from the Bern (Switzerland) blood donation center after obtaining the proper informed consent from healthy human donors. Blood samples were diluted 1:3 with sterile PBS and overlaid on Ficoll-Paque PLUS (GE Healthcare) using LeukoSep conical centrifuge tubes (Greiner Bio-One). Peripheral blood mononuclear cells (PBMCs) were harvested after separation for 30 minutes at 400 x *g* without braking. Successive centrifugation steps were performed to wash the mononuclear cell fraction and remove residual neutrophils and granulocytes. Total B cells were isolated from PBMCs by negative selection with the EasySep Human B Cell Enrichment Kit (STEMCELL Technologies) according to the manufacturer’s instructions. The following fluorescently labeled antibodies were used to stain the enriched B-cell fraction prior to sorting by flow cytometry: anti-CD3-APC/Cy7 (clone HIT3a BioLegend # 300318), anti-CD14-APC/Cy7 (clone HCD14 BioLegend #325620), anti-CD16-APC/Cy7 (clone 3G8 BioLegend #302018), anti-CD19-BV785 (clone HIB19, BioLegend #302240), anti-CD20-BV650 (clone 2H7, BioLegend #302336), anti-CD27-V450 (clone M-T271, BD Horizon #560448), anti-CD24-BV510 (clone ML5, BioLegend #311126), anti-CD38-PC5 (clone LS198-4-3, Beckman Coulter #A07780), anti-IgD-PEcy7 (clone IA6-2, BioLegend #348210), anti-IgG-Alexa Fluor 647 (Jackson ImmunoResearch #109-606-003) anti-IgA-Alexa Fluor 488 (Jackson ImmunoResearch #109-549-011), anti-IgM-PE (clone SA-DA4, eBioscience #12-9998). Cell sorting was performed on a BD FACS Aria III following the gating strategy depicted in Figure 4A. After sorting, isolated fractions were centrifuged 5 minutes at 300 x *g*, the supernatants were aspirated, and the cell pellets were re-suspended in 1 ml PBS. Recovered cells were hand-counted using a Neubauer hemocytometer, and aliquots containing equal numbers of cells were prepared from the cellular suspension. These aliquots were centrifuged, and the supernatant aspirated. Cell pellets were lysed directly in 200 µl TRI Reagent (Sigma), allowed to dissociate for 5 minutes at room temperature, then frozen on dry ice prior to storage at -80°C.

### RNA isolation from sorted B-cell populations

Immediately prior to use, Phase Lock Gel (PLG) tubes were pelleted at 12000 - 16000 x g in a microcentrifuge for 20 to 30 seconds. Each TRIzol aliquot was then thawed on ice. After thawing and an incubation time of 5 minutes at room temperature, 1 mL of the TRIzol homogenate was transferred to the phase lock tube. 0.2 mL chloroform was added and the tube was shaken vigorously by hand (∼15 seconds). After an incubation time of 3 minutes, the phase lock tube was centrifuged at 12’000 x g during 15 minutes at 4 °C. The resulting upper aqueous phase was transferred to a fresh Eppendorf tube, an equal volume of 70% ethanol was added and the solution was purified on a PureLink RNA column according to the manufacturer’s instructions (Life Technologies). Finally, RNA was eluted in 25 µl nuclease-free water.

### Library preparation and NGS on Illumina’s MiSeq

First-strand cDNA synthesis was carried out using Maxima reverse transcriptase (Life Technologies). The protocol for one reaction is as follows: A 29 µl reaction mix was prepared using up to 2 µg of RNA together with 40 pmol of the respective gene-specific reverse primer (for IgG sequences, 5’-RID-GTTCTGGGAAGTAGTCCTTGACCAG-3’ (IgH_10r); for IgM, 5’-RID-ACGAGGGGGAAAAGGGTTGG-3’ (CH1_1r)), 2 µl dNTP (10 mM each) and the required amount of nuclease-free water. This mix is incubated for 5 minutes at 65 °C. A master mix of 8 µl 5x RT buffer, 1 µl of (20 U) RiboLock and 2 µl of Maxima reverse transcriptase is then prepared and added to the reaction mix. Finally, the mix is incubated for 30 minutes at 50°C and the reaction is terminated by incubating at 85°C for 5 minutes.

The obtained cDNA is then cleaned with a left-sided SPRI-Select bead clean up (0.8x) according to the manufacturer’s instructions (Beckman coulter) and subsequently measured by ddPCR.

Up to 135,000 cDNA transcripts were then pooled together with 12,500 spike-in transcripts. Multiplex-PCR was performed using an equimolar pool of the forward primer mix (Figure 2B) and the reverse primer (TAK_423) targeting the overhang introduced during cDNA synthesis. Due to low cDNA yields, the first PCR was carried out for 20 cycles and the following cycling protocol: 2 min at 95 °C; 20 cycles of 98°C for 20s; 60°C for 50; 72°C for 1min; 72°C and then holding at 4°C. PCR reactions were prepared using 15 µl of Kapa HiFi HotStart ReadyMix (KAPA Biosystems), 50 pmol of the forward mix and the reverse primer and 9 µl of the cDNA mix. After the first PCR, we again performed a left-sided bead clean-up (0.8x) and measured the PCR product concentration using ddPCR. We use 800,000 transcripts from PCR 1 as input into the adapter extension PCR. For this PCR 25 µl of Kapa HiFI Hotstart ReadyMix was combined with 25 pmol of the forward primer (TAK_424) and 25 pmol of the index primer (TAK_531) as well as the diluted PCR product. Finally, the reaction volume was adjusted to 50 µl by the addition of nuclease-free water. Thermocycling was performed as follows: 95°C for 5 min; 23 cycles of 98°C for 20 s, 65°C for 15 s, 72°C for 15 s; 72°C for 5 min; and 4°C indefinitely. Following second-step adapter extension PCR, reactions were cleaned using a double-sided SPRIselect bead cleanup process (0.5x to 0.8x), with an additional ethanol wash and elution in TE buffer.

Libraries were then quantified by capillary electrophoresis (Fragment analyzer, Agilent). After quantitation, libraries were pooled accordingly and sequenced on a MiSeq System (Illumina) with the paired-end 2×300bp kit.

### Bioinformatic pipeline

Paired-end fastq files were merged using PandaSeq (40). Afterwards, sequences were filtered for quality and length using the FASTX toolkit (http://hannonlab.cshl.edu/fastx_toolkit/). After the quality trim, sequences were processed with a custom Python script that performed error correction by consensus building on our sequences and RIDs. In order to utilize as many sequencing reads as possible, we required UIDs to have at least 3 reads, but did not remove sequences that only had one UID group mapping to them. VDJ annotation and frequency calculation was then performed by our in-house aligner (18) which was updated with the human reference database downloaded from IMGT. The complete error-correction and alignment pipeline is available under https://gitlab.ethz.ch/reddy/MAF.

### Statistical analysis

All statistical and computational analyses following the alignment step were performed in R. Details about specific tests that were used can be found in the results section and in the figure legends. Scripts are available upon request.

## Data availability

In adherence to the data sharing recommendations of the AIRR community our data is publically available in the following repositories: BioProject, BioSample, SRA and GenBank and can be accessed with the accession number PRJNA430091 (BioProject). The exact data processing steps, including software tools and version numbers can be found on zonodo.org under the following doi: 10.5281/zenodo.1201416.

Likewise, the designed spike-in sequences are also stored on GenBank (Accession number MG785894-MG785978).

## Author contributions

J.M.L., S.F., E.T., and S.T.R. designed experiments. V.C. performed B-cell enrichment, sorting, and mRNA extraction. M.I. and S.F. prepared IgH libraries. J.M.L. designed primer sequences. J.M.L. and A.Z. designed antibody spike-ins. A.Z., S.M., and M.I. conducted preliminary experiments. E.M. analysed preliminary data. S.F. was responsible for the bioinformatics pipeline. J.M.L. and S.F. analysed the final data and prepared figures. J.M.L., S.F., E.T., and S.T.R. wrote the paper. All authors provided scientific guidance.

## Acknowledgements

We would like to acknowledge the Genomics Facility Basel of ETH Zurich for Illumina sequencing support, in particular E. Burcklen, K. Eschbach and C. Beisel. We also want to thank H. Ruscheweyh for bioinformatic code support.

