## Supplementary Materials for "Synthetic standards combined with error and bias correction improves the accuracy and quantitative resolution of antibody repertoire sequencing in human naïve and memory B cells"

### Supplementary Material

| CDR3 amino acid sequence | CDR3 length | Variable (V) gene segment | Joining (J) gene segment | VH family | JH | IgG subclass | total mismatches | FR1 mismatches | CDR1 mismatches | FR2 mismatches | CDR2 mismatches | FR3 mismatches | spike-in pool contribution (%) |
|---|---|---|---|---|---|---|---|---|---|---|---|---|---|
| ARYANDREASY | 11 | IGHV3-23*01 | IGHJ4*01 | VH3 | JH4 | IgG3 | 0 | 0 | 0 | 0 | 0 | 0 | 7.00 |
| ARCAFELATTE | 11 | IGHV3-23*01 | IGHJ6*01 | VH3 | JH6 | IgG2 | 9 | 0 | 4 | 0 | 5 | 0 | 0.15 |
| ARSIMNFRIED | 11 | IGHV3-23*01 | IGHJ1*01 | VH3 | JH1 | IgG3 | 9 | 0 | 4 | 1 | 3 | 1 | 1.11 |
| ARDRSAIREDDY | 12 | IGHV4-34*01 | IGHJ4*01 | VH4 | JH4 | IgG4 | 0 | 0 | 0 | 0 | 0 | 0 | 0.21 |
| ARCRAIGVENTER | 13 | IGHV4-34*12 | IGHJ6*01 | VH4 | JH6 | IgG3 | 6 | 0 | 2 | 0 | 3 | 1 | 5.92 |
| ARCHEETAHTEN | 12 | IGHV4-34*12 | IGHJ3*01 | VH4 | JH3 | IgG2 | 8 | 0 | 4 | 0 | 4 | 0 | 0.57 |
| ARTHERESAPESCH | 14 | IGHV1-69*06 | IGHJ4*01 | VH1 | JH4 | IgG3 | 0 | 0 | 0 | 0 | 0 | 0 | 0.18 |
| ARSSQIRREL | 10 | IGHV1-69*06 | IGHJ6*01 | VH1 | JH6 | IgG4 | 5 | 0 | 1 | 1 | 3 | 0 | 0.80 |
| ARTARIKKHAN | 11 | IGHV1-69*06 | IGHJ1*01 | VH1 | JH1 | IgG3 | 10 | 0 | 4 | 0 | 5 | 1 | 3.04 |
| ARCAMELLIATREE | 14 | IGHV1-18*01 | IGHJ4*01 | VH1 | JH4 | IgG2 | 0 | 0 | 0 | 0 | 0 | 0 | 0.25 |
| ARCRISTINAPA | 12 | IGHV1-18*01 | IGHJ6*01 | VH1 | JH6 | IgG3 | 10 | 0 | 4 | 3 | 2 | 1 | 0.68 |
| ARTETRACYCLINE | 14 | IGHV1-18*01 | IGHJ3*01 | VH1 | JH3 | IgG4 | 12 | 0 | 4 | 4 | 4 | 0 | 2.57 |
| ARNICEECHIDNA | 13 | IGHV4-61*08 | IGHJ4*01 | VH4 | JH4 | IgG3 | 0 | 0 | 0 | 0 | 0 | 0 | 0.21 |
| ARGIANTPANDA | 12 | IGHV4-61*08 | IGHJ6*01 | VH4 | JH6 | IgG2 | 7 | 0 | 3 | 0 | 4 | 0 | 0.94 |
| ARGIRAFFEAFFE | 13 | IGHV4-61*08 | IGHJ1*01 | VH4 | JH1 | IgG3 | 15 | 0 | 4 | 3 | 8 | 0 | 3.04 |
| ARIMPALAPALA | 12 | IGHV4-59*01 | IGHJ5*01 | VH4 | JH5 | IgG4 | 0 | 0 | 0 | 0 | 0 | 0 | 0.25 |
| ARRATSEATMICE | 13 | IGHV4-59*01 | IGHJ6*01 | VH4 | JH6 | IgG3 | 10 | 0 | 4 | 0 | 4 | 2 | 0.48 |
| ARRATTLESNAKE | 13 | IGHV4-59*01 | IGHJ3*01 | VH4 | JH3 | IgG2 | 10 | 1 | 5 | 0 | 3 | 1 | 2.17 |
| ARREINDEER | 10 | IGHV3-30*03 | IGHJ4*01 | VH3 | JH4 | IgG3 | 0 | 0 | 0 | 0 | 0 | 0 | 0.18 |
| ARSALAMANDER | 12 | IGHV3-30*03 | IGHJ6*01 | VH3 | JH6 | IgG4 | 9 | 0 | 3 | 2 | 4 | 0 | 1.32 |
| ARTIGERSEVEN | 12 | IGHV3-30*03 | IGHJ1*01 | VH3 | JH1 | IgG3 | 12 | 0 | 4 | 0 | 5 | 3 | 2.17 |
| ARTREESHREW | 11 | IGHV4-39*01 | IGHJ4*01 | VH4 | JH4 | IgG2 | 0 | 0 | 0 | 0 | 0 | 0 | 0.29 |
| ARMANHATTAN | 11 | IGHV4-39*01 | IGHJ6*01 | VH4 | JH6 | IgG3 | 5 | 0 | 1 | 0 | 4 | 0 | 0.94 |
| ARIRISHCAFE | 11 | IGHV4-39*01 | IGHJ3*01 | VH4 | JH3 | IgG4 | 11 | 2 | 3 | 2 | 4 | 0 | 1.84 |
| ARNEWWHISKEY | 12 | IGHV3-48*02 | IGHJ4*01 | VH3 | JH4 | IgG3 | 0 | 0 | 0 | 0 | 0 | 0 | 0.35 |
| ARSWEETWINE | 11 | IGHV3-48*02 | IGHJ6*01 | VH3 | JH6 | IgG2 | 7 | 0 | 3 | 0 | 4 | 0 | 1.56 |
| ARGINMADRAS | 11 | IGHV3-48*02 | IGHJ2*01 | VH3 | JH2 | IgG3 | 11 | 0 | 4 | 1 | 6 | 0 | 1.84 |
| ARAPPLETINIS | 12 | IGHV1-3*02 | IGHJ4*01 | VH1 | JH4 | IgG4 | 0 | 0 | 0 | 0 | 0 | 0 | 0.35 |
| ARCAIPIRINHA | 12 | IGHV1-3*02 | IGHJ4*01 | VH1 | JH4 | IgG3 | 6 | 0 | 2 | 0 | 4 | 0 | 1.32 |
| ARNECTARINE | 11 | IGHV3-21*01 | IGHJ1*01 | VH3 | JH1 | IgG2 | 0 | 0 | 0 | 0 | 0 | 0 | 0.41 |
| ARPHYSALIS | 10 | IGHV3-21*01 | IGHJ2*01 | VH3 | JH2 | IgG3 | 10 | 0 | 5 | 0 | 3 | 2 | 0.80 |
| ARAAMARANTH | 11 | IGHV1-2*02 | IGHJ6*01 | VH1 | JH6 | IgG4 | 0 | 0 | 0 | 0 | 0 | 0 | 0.57 |
| ARGRAPESALAD | 12 | IGHV1-2*02 | IGHJ4*01 | VH1 | JH4 | IgG3 | 11 | 0 | 7 | 0 | 3 | 1 | 0.21 |
| ARFIDDLEHEAD | 12 | IGHV4-31*02 | IGHJ4*01 | VH4 | JH4 | IgG2 | 0 | 0 | 0 | 0 | 0 | 0 | 0.48 |
| ARFRIEDGARLIC | 13 | IGHV4-31*02 | IGHJ1*01 | VH4 | JH1 | IgG3 | 8 | 0 | 2 | 0 | 6 | 0 | 0.15 |
| ARFATCHICKEN | 12 | IGHV3-33*01 | IGHJ2*01 | VH3 | JH2 | IgG4 | 0 | 0 | 0 | 0 | 0 | 0 | 0.48 |
| ARDRIEDSEAWEED | 14 | IGHV3-33*01 | IGHJ6*01 | VH3 | JH6 | IgG3 | 8 | 0 | 3 | 0 | 1 | 4 | 1.56 |
| ARSWISSCHARD | 12 | IGHV5-51*01 | IGHJ4*01 | VH5 | JH4 | IgG2 | 0 | 0 | 0 | 0 | 0 | 0 | 0.29 |
| ARWATERCRESS | 12 | IGHV5-51*01 | IGHJ4*01 | VH5 | JH4 | IgG3 | 2 | 0 | 2 | 0 | 0 | 0 | 1.11 |
| AREGGPLANT | 10 | IGHV1-46*01 | IGHJ1*01 | VH1 | JH1 | IgG4 | 0 | 0 | 0 | 0 | 0 | 0 | 0.41 |
| ARARGENTINA | 11 | IGHV1-46*01 | IGHJ6*01 | VH1 | JH6 | IgG3 | 5 | 0 | 3 | 0 | 1 | 1 | 0.18 |
| ARAEINSTEIN | 11 | IGHV3-7*01 | IGHJ4*01 | VH3 | JH4 | IgG2 | 0 | 0 | 0 | 0 | 0 | 0 | 0.29 |
| ARKASACHSTAN | 12 | IGHV3-7*01 | IGHJ4*01 | VH3 | JH4 | IgG3 | 8 | 0 | 4 | 0 | 4 | 0 | 0.94 |
| ARAPICCARD | 10 | IGHV4-38-2*02 | IGHJ1*01 | VH4 | JH1 | IgG2 | 0 | 0 | 0 | 0 | 0 | 0 | 0.25 |
| ARNETHERLANDS | 13 | IGHV4-38-2*02 | IGHJ6*01 | VH4 | JH6 | IgG3 | 6 | 0 | 1 | 1 | 4 | 0 | 0.80 |
| ARSELFIESTICK | 13 | IGHV3-11*01 | IGHJ4*01 | VH3 | JH4 | IgG2 | 0 | 0 | 0 | 0 | 0 | 0 | 0.41 |
| ARPEKINGENTE | 12 | IGHV3-11*01 | IGHJ3*01 | VH3 | JH3 | IgG3 | 6 | 0 | 3 | 1 | 2 | 0 | 1.32 |
| ARCHAIMASALA | 12 | IGHV1-8*01 | IGHJ1*01 | VH1 | JH1 | IgG2 | 2 | 0 | 0 | 2 | 0 | 0 | 0.21 |
| ARGINGERALE | 11 | IGHV1-8*01 | IGHJ6*01 | VH1 | JH6 | IgG3 | 6 | 1 | 0 | 4 | 1 | 0 | 0.68 |
| ARGREATREDMEAT | 14 | IGHV3-66*03 | IGHJ5*01 | VH3 | JH5 | IgG2 | 0 | 0 | 0 | 0 | 0 | 0 | 0.68 |
| ARRACLETTE | 10 | IGHV3-66*03 | IGHJ3*01 | VH3 | JH3 | IgG3 | 15 | 0 | 3 | 2 | 6 | 4 | 0.25 |
| ARAMYSMART | 10 | IGHV3-30*04 | IGHJ1*01 | VH3 | JH1 | IgG2 | 0 | 0 | 0 | 0 | 0 | 0 | 0.29 |
| ARENDLESSWAY | 12 | IGHV3-30*04 | IGHJ6*01 | VH3 | JH6 | IgG3 | 11 | 1 | 4 | 4 | 2 | 0 | 0.80 |
| ARLANADELREY | 12 | IGHV2-5*01 | IGHJ4*01 | VH2 | JH4 | IgG2 | 0 | 0 | 0 | 0 | 0 | 0 | 4.24 |
| ARCDELEVINGNE | 13 | IGHV2-5*01 | IGHJ3*01 | VH2 | JH3 | IgG3 | 3 | 0 | 3 | 0 | 0 | 0 | 1.11 |
| ARTHEISLAND | 11 | IGHV3-64*02 | IGHJ2*01 | VH3 | JH2 | IgG3 | 0 | 0 | 0 | 0 | 0 | 0 | 3.59 |
| ARRFEDERER | 10 | IGHV3-64*02 | IGHJ6*01 | VH3 | JH6 | IgG3 | 8 | 0 | 4 | 1 | 3 | 0 | 0.57 |
| ARVANILLACAKE | 13 | IGHV4-4*07 | IGHJ4*01 | VH4 | JH4 | IgG3 | 0 | 0 | 0 | 0 | 0 | 0 | 0.35 |
| ARWALLSTREET | 12 | IGHV4-4*07 | IGHJ3*01 | VH4 | JH3 | IgG3 | 7 | 0 | 2 | 0 | 5 | 0 | 1.11 |
| ARGRAYALDER | 11 | IGHV3-15*01 | IGHJ2*01 | VH3 | JH2 | IgG3 | 0 | 0 | 0 | 0 | 0 | 0 | 1.84 |
| ARCHERRYPIE | 11 | IGHV3-15*01 | IGHJ6*01 | VH3 | JH6 | IgG3 | 4 | 0 | 0 | 2 | 2 | 0 | 0.25 |
| ARSPICYSHRIMP | 13 | IGHV2-26*01 | IGHJ5*01 | VH2 | JH5 | IgG3 | 0 | 0 | 0 | 0 | 0 | 0 | 0.68 |
| ARFRESHTHYME | 12 | IGHV2-26*01 | IGHJ3*01 | VH2 | JH3 | IgG3 | 6 | 0 | 4 | 0 | 2 | 0 | 0.41 |
| ARTRAVELLING | 12 | IGHV3-74*01 | IGHJ2*01 | VH3 | JH2 | IgG3 | 0 | 0 | 0 | 0 | 0 | 0 | 0.94 |
| ARFMAGELLAN | 11 | IGHV3-74*01 | IGHJ6*01 | VH3 | JH6 | IgG3 | 7 | 0 | 1 | 2 | 4 | 0 | 0.35 |
| ARSWISSNESS | 11 | IGHV2-70*13 | IGHJ1*01 | VH2 | JH1 | IgG3 | 0 | 0 | 0 | 0 | 0 | 0 | 0.15 |
| ARMILKSHAKE | 11 | IGHV4-30-4*01 | IGHJ2*01 | VH4 | JH2 | IgG3 | 0 | 0 | 0 | 0 | 0 | 0 | 0.48 |
| ARGREENPEPPER | 13 | IGHV7-4-1*02 | IGHJ4*01 | VH7 | JH4 | IgG3 | 0 | 0 | 0 | 0 | 0 | 0 | 0.57 |
| ARANGELFALLS | 12 | IGHV3-53*01 | IGHJ6*01 | VH3 | JH6 | IgG3 | 0 | 0 | 0 | 0 | 0 | 0 | 0.29 |
| ARDEATHVALLEY | 13 | IGHV6-1*01 | IGHJ1*01 | VH6 | JH1 | IgG3 | 0 | 0 | 0 | 0 | 0 | 0 | 0.41 |
| ARVARANASI | 10 | IGHV4-30-2*03 | IGHJ3*02 | VH4 | JH3 | IgG3 | 0 | 0 | 0 | 0 | 0 | 0 | 0.35 |
| ARSTARWARS | 10 | IGHV4-28*01 | IGHJ4*01 | VH4 | JH4 | IgG3 | 0 | 0 | 0 | 0 | 0 | 0 | 4.24 |
| ARFISHNCHIPS | 12 | IGHV3-9*01 | IGHJ6*01 | VH3 | JH6 | IgG3 | 0 | 0 | 0 | 0 | 0 | 0 | 5.92 |
| ARITSATEST | 10 | IGHV3-20*01 | IGHJ2*01 | VH3 | JH2 | IgG3 | 0 | 0 | 0 | 0 | 0 | 0 | 2.57 |
| ARHERAKLES | 10 | IGHV1-24*01 | IGHJ3*01 | VH1 | JH3 | IgG3 | 0 | 0 | 0 | 0 | 0 | 0 | 4.24 |
| ARWILLIAMKELT | 13 | IGHV3-49*03 | IGHJ5*01 | VH3 | JH5 | IgG3 | 0 | 0 | 0 | 0 | 0 | 0 | 3.04 |
| ARMINERVATHECAT | 15 | IGHV1-69-2*01 | IGHJ6*01 | VH1 | JH6 | IgG3 | 2 | 0 | 0 | 0 | 2 | 0 | 1.56 |
| ARTARDIGRADA | 9 | IGHV5-10-1*02 | IGHJ2*01 | VH5 | JH2 | IgG3 | 0 | 0 | 0 | 0 | 0 | 0 | 1.32 |
| ARRFRANKLIN | 11 | IGHV1-58*02 | IGHJ3*01 | VH1 | JH3 | IgG3 | 0 | 0 | 0 | 0 | 0 | 0 | 0.80 |
| ARSELACHII | 10 | IGHV3-72*01 | IGHJ4*01 | VH3 | JH4 | IgG3 | 0 | 0 | 0 | 0 | 0 | 0 | 0.57 |
| ARTEHRANINIRAN | 14 | IGHV3-73*02 | IGHJ6*01 | VH3 | JH6 | IgG3 | 0 | 0 | 0 | 0 | 0 | 0 | 0.18 |
| ARDAMAVAND | 10 | IGHV3-13*01 | IGHJ2*01 | VH3 | JH2 | IgG3 | 0 | 0 | 0 | 0 | 0 | 0 | 0.21 |
| ARNEMERTEA | 10 | IGHV1-45*02 | IGHJ4*01 | VH1 | JH4 | IgG3 | 0 | 0 | 0 | 0 | 0 | 0 | 0.48 |
| ARKANAMYCIN | 11 | IGHV3-43*01 | IGHJ4*01 | VH3 | JH4 | IgG3 | 0 | 0 | 0 | 0 | 0 | 0 | 0.68 |
| ARTRAGGIAI | 10 | IGHV7-81*01 | IGHJ6*01 | VH7 | JH6 | IgG3 | 0 | 0 | 0 | 0 | 0 | 0 | 0.48 |

**Supplementary table 1.** Molecular characteristics of 85 synthetic human IgH gene standards ('spike-ins') used in this study. See Figure 1A for spike-in construction schematic.

| Sample | Total RNA extracted (ng/uL) | Total RNA (total amount) | Average number of cDNA transcripts per uL | Total number of cDNA transcripts | Sequencing depth (number of reads prior to pre-procssing) | Number of reads after pre-processing |
|---|---|---|---|---|---|---|
| IgM1_D1 | 6.6 | 132 | 5'109 | 62'581 | 676'557 | 403'271 |
| IgM2_D1 | 59.1 | 1182 | 4'691 | 57'469 | 781'452 | 508'998 |
| IgM3_D1 | 10.1 | 202 | 3'503 | 42'916 | 2'236'338 | 1'039'415 |
| IgM4_D1 | 30.6 | 612 | 405 | 4'955 | 792'330 | 437'560 |
| IgM5_D1 | 19 | 380 | 3'293 | 40'333 | 1'519'422 | 529'933 |
| IgM1_D2 | 3.7 | 74 | 3'390 | 41'528 | 1'925'305 | 1'111'250 |
| IgM2_D2 | 53.3 | 1066 | 4'330 | 53'043 | 673'482 | 379'811 |
| IgM3_D2 | 34.4 | 688 | 2'048 | 25'088 | 992'456 | 488'230 |
| IgM4_D2 | 2.2 | 44 | 3'075 | 37'669 | 981'274 | 452'817 |
| IgM5_D2 | 47.4 | 948 | 3'855 | 47'220 | 2'279'624 | 440'083 |
| IgM1_D3 | 7.7 | 154 | 4'631 | 56'734 | 545'947 | 333'492 |
| IgM2_D3 | 29.5 | 590 | 3'568 | 43'702 | 1'448'340 | 636'432 |
| IgM3_D3 | 2.6 | 52 | 2'860 | 35'035 | 649'626 | 407'914 |
| IgM4_D3 | 7.2 | 144 | 3'567 | 43'696 | 986'419 | 529'860 |
| IgG1_D1 | 61.2 | 1224 | 1'277 | 15'643 | 4'282'524 | 1'005'440 |
| IgG2_D1 | 8.7 | 174 | 1'289 | 15'784 | 2'513'370 | 2'099'114 |
| IgG3_D1 | 12.1 | 242 | 1'549 | 18'969 | 720'804 | 635'650 |
| IgG4_D1 | 9.4 | 188 | 1'577 | 19'312 | 1'057'258 | 919'252 |
| IgG5_D1 | 30.7 | 614 | 2'884 | 35'325 | 1'497'219 | 1'306'234 |
| IgG1_D2 | 52.4 | 1048 | 421 | 5'157 | 1'049'571 | 913'558 |
| IgG2_D2 | 3.7 | 74 | 181 | 2'211 | 959'085 | 700'808 |
| IgG3_D2 | 8.5 | 170 | 268 | 3'277 | 910'336 | 793'379 |
| IgG4_D2 | 6.5 | 130 | 214 | 2'622 | 747'740 | 662'657 |
| IgG5_D2 | 5.6 | 112 | 386 | 4'722 | 998'038 | 826'427 |
| IgG1_D3 | 7 | 140 | 1'552 | 19'012 | 604'835 | 490'991 |
| IgG2_D3 | 5.4 | 108 | 683 | 8'367 | 804'122 | 658'452 |
| IgG3_D3 | 5.9 | 118 | 972 | 11'901 | 756'943 | 649'224 |
| IgG4_D3 | 6.9 | 138 | 1'639 | 20'072 | 2'724'771 | 1'179'968 |

**Supplementary table 2**. Overview over our experimental results

| Sample | Aligned reads (consensus build) | Raw CDR3 variants (AA) | Raw CDR3 variants w.o. singletons (AA) | Raw total variants (whole VDJ, nt) | MAF corrected CDR3 variants (AA) | MAF corrected clonotypes (Same V/J Gene, same | MAF corrected variants (whole VDJ, nt) | Donor |
|---|---|---|---|---|---|---|---|---|
| IgM1_D1 | 201'889 | 40'584 | 19'815 | 109'404 | 14'891 | 14'629 | 18'120 | D1 |
| IgM2_D1 | 260'867 | 37'987 | 14'952 | 129'770 | 11'257 | 11'088 | 13'629 | D1 |
| IgM3_D1 | 393'067 | 47'447 | 17'357 | 155'913 | 11'759 | 11'639 | 14'106 | D1 |
| IgM4_D1 | 203'127 | 46'001 | 21'116 | 128'936 | 15'139 | 14'919 | 18'001 | D1 |
| IgM5_D1 | 175'215 | 32'357 | 14'870 | 96'251 | 11'375 | 11'185 | 14'088 | D1 |
| IgM1_D2 | 534'227 | 65'874 | 24'260 | 234'442 | 18'087 | 17'646 | 23'645 | D2 |
| IgM2_D2 | 214'469 | 37'837 | 16'146 | 117'594 | 12'255 | 12'055 | 14'567 | D2 |
| IgM3_D2 | 244'601 | 35'706 | 14'292 | 120'009 | 10'643 | 10'439 | 13'289 | D2 |
| IgM4_D2 | 204'029 | 37'669 | 16'697 | 111'390 | 12'491 | 12'269 | 15'254 | D2 |
| IgM5_D2 | 135'195 | 26'236 | 12'557 | 66'889 | 9'397 | 9'292 | 10'797 | D2 |
| IgM1_D3 | 179'806 | 32'421 | 12'987 | 97'529 | 9'960 | 9'808 | 12'518 | D3 |
| IgM2_D3 | 285'749 | 42'979 | 18'424 | 138'162 | 14'098 | 13'866 | 17'610 | D3 |
| IgM3_D3 | 197'051 | 33'712 | 15'530 | 101'684 | 11'748 | 11'567 | 14'455 | D3 |
| IgM4_D3 | 248'200 | 31'542 | 12'514 | 103'246 | 9'693 | 9'582 | 11'799 | D3 |
| IgG1_D1 | 207'903 | 32'401 | 9'126 | 82'538 | 7'161 | 6'756 | 8'963 | D1 |
| IgG2_D1 | 1'484'045 | 159'576 | 36'883 | 520'865 | 8'549 | 7'744 | 13'393 | D1 |
| IgG3_D1 | 489'704 | 83'795 | 19'308 | 246'780 | 10'986 | 9'957 | 15'014 | D1 |
| IgG4_D1 | 712'466 | 105'073 | 24'949 | 322'613 | 11'320 | 10'286 | 17'581 | D1 |
| IgG5_D1 | 1'025'422 | 162'369 | 38'540 | 489'854 | 18'690 | 16'575 | 28'718 | D1 |
| IgG1_D2 | 644'950 | 56'789 | 13'373 | 199'164 | 3'233 | 3'011 | 4'954 | D2 |
| IgG2_D2 | 429'902 | 24'874 | 6'368 | 80'817 | 1'650 | 1'456 | 2'767 | D2 |
| IgG3_D2 | 541'960 | 38'086 | 8'957 | 134'139 | 1'965 | 1'849 | 2'895 | D2 |
| IgG4_D2 | 472'428 | 25'158 | 5'943 | 83'147 | 1'398 | 1'288 | 2'043 | D2 |
| IgG5_D2 | 566'689 | 40'047 | 9'394 | 132'147 | 2'501 | 2'323 | 3'603 | D2 |
| IgG1_D3 | 367'556 | 74'198 | 16'725 | 207'180 | 10'665 | 9'808 | 14'843 | D3 |
| IgG2_D3 | 485'151 | 55'285 | 12'829 | 182'680 | 4'559 | 4'266 | 6'327 | D3 |
| IgG3_D3 | 502'248 | 60'245 | 14'155 | 206'903 | 5'430 | 5'082 | 7'687 | D3 |
| IgG4_D3 | 693'622 | 96'133 | 22'466 | 325'211 | 9'239 | 8'556 | 14'028 | D3 |

| Primer | Sequence | Notes |
|---|---|---|
| IgG_1r | TTGGCACCCGAGAATTCCACTGHHHHHACAHHHHHACAHHHHNATTGTTCTGGGAAGTAGTCCTTGACCAG | Red part indicates primer binding site, black part contains unique identifier and overhang |
| IgM_1r | TTGGCACCCGAGAATTCCACTGHHHHHACAHHHHHACAHHHHNATTACGAGGGGGAAAAGGGTTGG | see above |
| VH1a | CGTTCAGAGTTCTACAGTCCGACGATCHHHHACHHHHACHHHNGCAGGCAGCTGGTGCAGTCTGGGG | see above |
| VH1b | CGTTCAGAGTTCTACAGTCCGACGATCHHHHACHHHHACHHHNGCAGTCAGCTGGTGCAGTCTGGAG | see above |
| VH1c | CGTTCAGAGTTCTACAGTCCGACGATCHHHHACHHHHACHHHNGCAGCCAGCTTGTGCAGTCTGGGG | see above |
| VH1d | CGTTCAGAGTTCTACAGTCCGACGATCHHHHACHHHHACHHHNGCAGGCAGCTGGTGCAGTCTGGGC | see above |
| VH2a | CGTTCAGAGTTCTACAGTCCGACGATCHHHHACHHHHACHHHNGCAGCCCAGGTCACCTTGAAGGAGTCTG | see above |
| VH2b | CGTTCAGAGTTCTACAGTCCGACGATCHHHHACHHHHACHHHNGCAGCCCAGATCACCTTGAAGGAGTCTG | see above |
| VH3a | CGTTCAGAGTTCTACAGTCCGACGATCHHHHACHHHHACHHHNGCAGTGTGAGGTGCAGCTGGTGGAGTC | see above |
| VH3b | CGTTCAGAGTTCTACAGTCCGACGATCHHHHACHHHHACHHHNGCAGTGTGAAGTGCAGCTGGTGGAGTC | see above |
| VH3c | CGTTCAGAGTTCTACAGTCCGACGATCHHHHACHHHHACHHHNGCAGTGTCAGGTGCAGCTGGTGGAGTC | see above |
| VH3d | CGTTCAGAGTTCTACAGTCCGACGATCHHHHACHHHHACHHHNGCAGTGTGAGGTGCAGCTGGTGGAGAC | see above |
| VH4a | CGTTCAGAGTTCTACAGTCCGACGATCHHHHACHHHHACHHHNGCAGTGCAGCTGCAGGAGTCGGG | see above |
| VH4b | CGTTCAGAGTTCTACAGTCCGACGATCHHHHACHHHHACHHHNGCAGTACAGCTGCAGGAGTCGGG | see above |
| VH4c | CGTTCAGAGTTCTACAGTCCGACGATCHHHHACHHHHACHHHNGCAGTGCAGCTGCAGGAGTCCGG | see above |
| VH4d | CGTTCAGAGTTCTACAGTCCGACGATCHHHHACHHHHACHHHNGCAGTGCAGCTACAGCAGTGGGG | see above |
| VH5 | CGTTCAGAGTTCTACAGTCCGACGATCHHHHACHHHHACHHHNGCAGGCAGCTGGTGCAGTCTGGAG | see above |
| VH6 | CGTTCAGAGTTCTACAGTCCGACGATCHHHHACHHHHACHHHNGCAGTACAGCTGCAGCAGTCAGG | see above |
| VH7 | CGTTCAGAGTTCTACAGTCCGACGATCHHHHACHHHHACHHHNGCAGTGCAGCTGGTGCAATCTGG | see above |
| PCR1_1r | ACTGGAGTTCCTTGGCACCCGAGAATTCCACT*G | '_*' indicates phosphorothioate bond |
| PCR2_f | AATGATACGGCGACCACCGAGATCTACACGTTCAGAGTTCTACAGTCCGACGAT*C | '_*' indicates phosphorothioate bond |
| PCR2_r | CAAGCAGAAGACGGCATACGAGATXXXXXXGTGACTGGAGTTCCTTGGCACCC | Red X indicates Illumina index |
| IgG_probe | /56-FAM/T+CTT+CCCC+C+TG+G/3IABkFQ/ | '+_' indicate locked ribonucleic acids, '/56'-FAM = 5' 6-FAM (Fluorescein), '/3IABkFQ/' = 3' Iowa Black FQ |
| IgM_probe | /56-FAM/C+CCC+AA+CC+C+TTT/3IABkFQ/ | '+_' indicate locked ribonucleic acids, '/56'-FAM = 5' 6-FAM (Fluorescein), '/3IABkFQ/' = 3' Iowa Black FQ |
| Spike_probe | /5HEX/CG+T C+T+G ACT +AGA +ACT +C/3IABkFQ/ | '+_' indicate locked ribonucleic acids, '/56'-FAM = 5' HEX (Hexachlorofluorescein),'/3IABkFQ/' = 3' Iowa Black FQ |
| ddPCR_f | GGTCACYGTCTCYTCAG | |
| ddPCR_r | TGGCACCCGAGAATTC | |

**Supplementary table 3.** Overview over all primers and probes used in this study

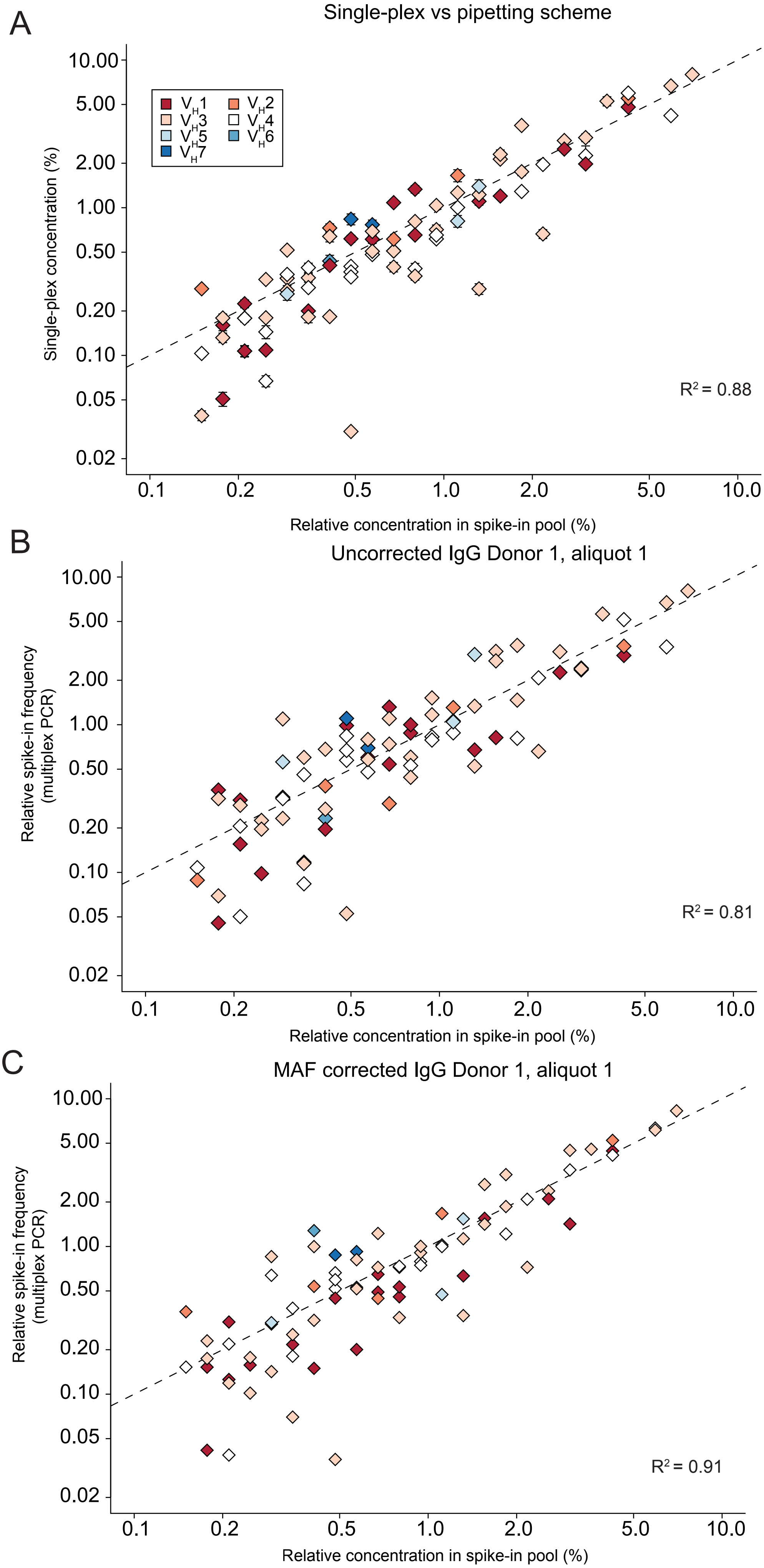


**Figure S1.** The sequencing bias of the multiplexed primer set was assessed by plotting the measured frequencies of each standard versus its actual, pipetted concentration in the pool. In an ideal case, the measured frequencies would fall onto the dashed line. The deviation from this line was used to calculate the R2 value, which decreases with greater deviation. (A) The upper panel shows the divergence of the measured frequencies obtained in the singleplex experiments versus pipetted concentration. Panels (B) and (C) show the measured frequencies and their deviations for one single experiment (IgG, Donor 1, Aliquot 1) before and after MAF correction.

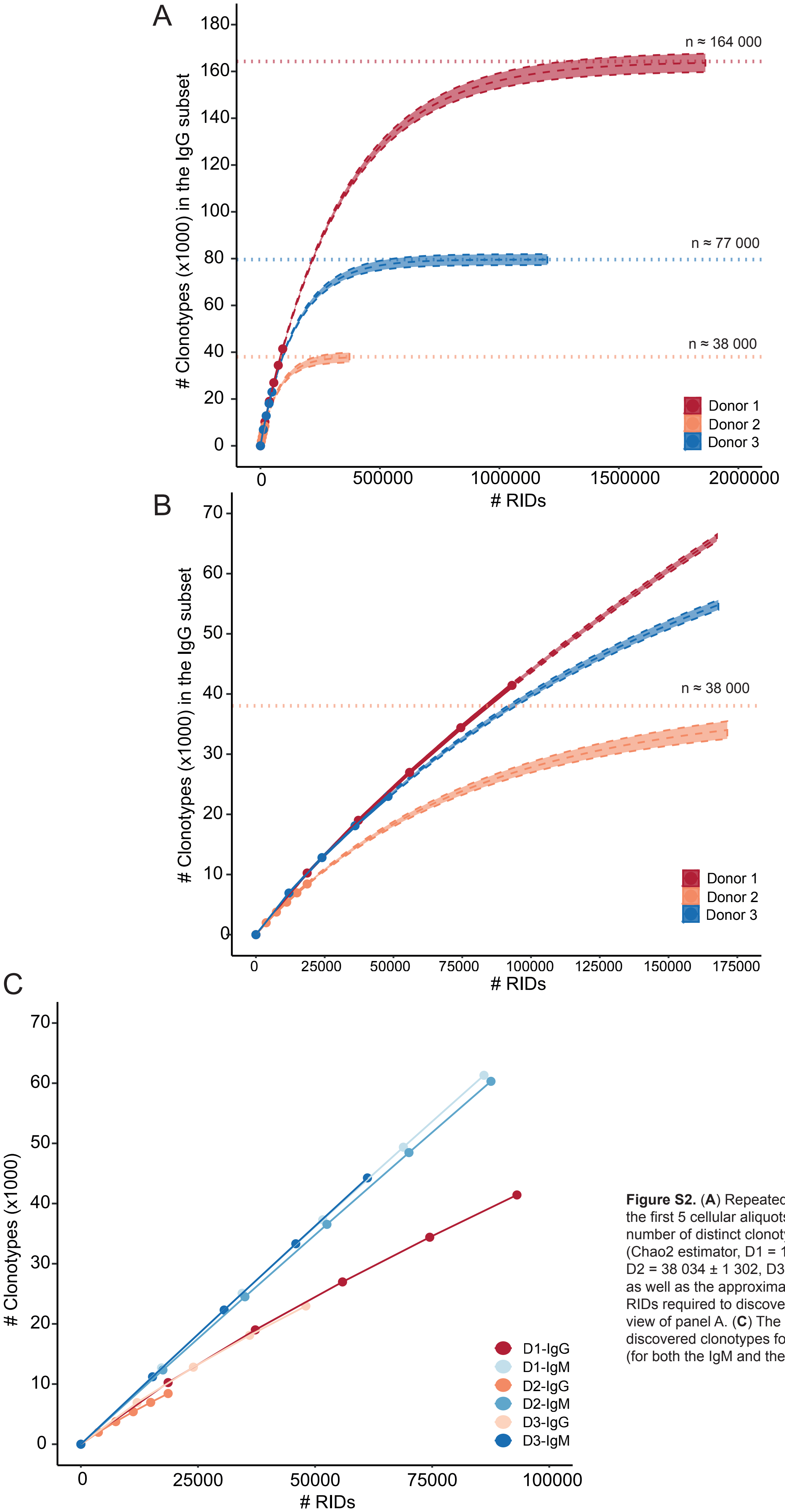


**Figure S2.** (**A**) Repeatedly observed sequences within the first 5 cellular aliquots allow determination of the number of distinct clonotypes in the IgG CD27+ subset (Chao2 estimator, D1 = 164 268 ± 2 365, D2 = 38 034 ± 1 302, D3 = 76 904 ± 1 409), as well as the approximate amount of cDNA transcripts / RIDs required to discover all clonotypes. (**B**) Expanded view of panel A. (**C**) The average number of newly discovered clonotypes for each additional replicate (for both the IgM and the IgG subset).

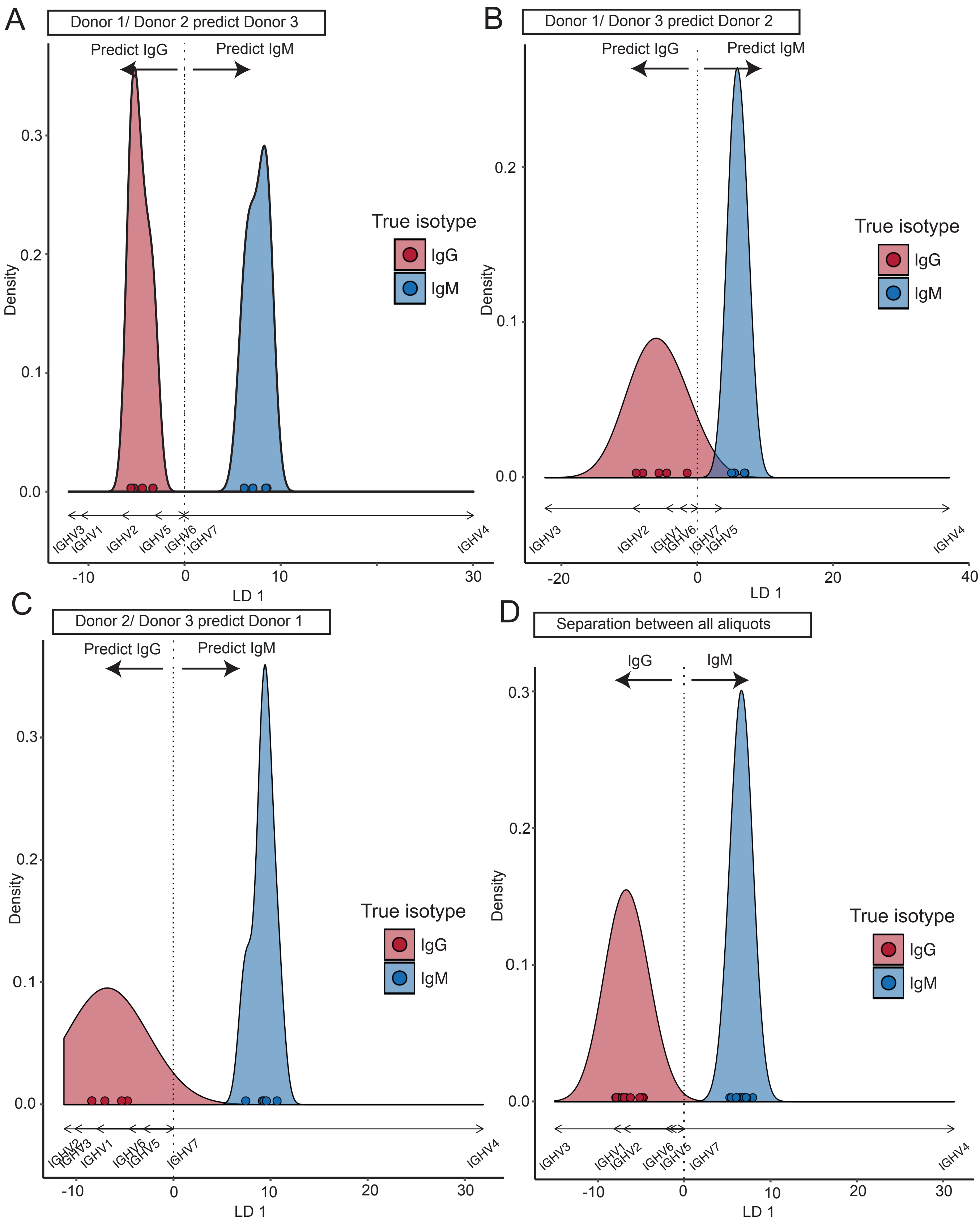


Fig. S3. **Linear discriminant analysis distinguishes CD27+ IgG repertoires and CD27- IgM repertoires based on their V-Gene family usage.** (**A**) All aliquots from donors 1 and 2 were used to fit an LDA classifier based on the centered log ratio transformed V-Gene family frequencies. Afterwards, the aliquots from donor 3 are projected to the fitted component axis. Positive and negative values predict IgM and IgG repertoires, respectively. Arrows below the plot indicate the contribution of each V-Gene family to the prediction. Colored dots show true class membership and their positions are smoothed using kernel desnity estimators. Panels (**B**-**D**) shows the same procedure for different splits of the data.

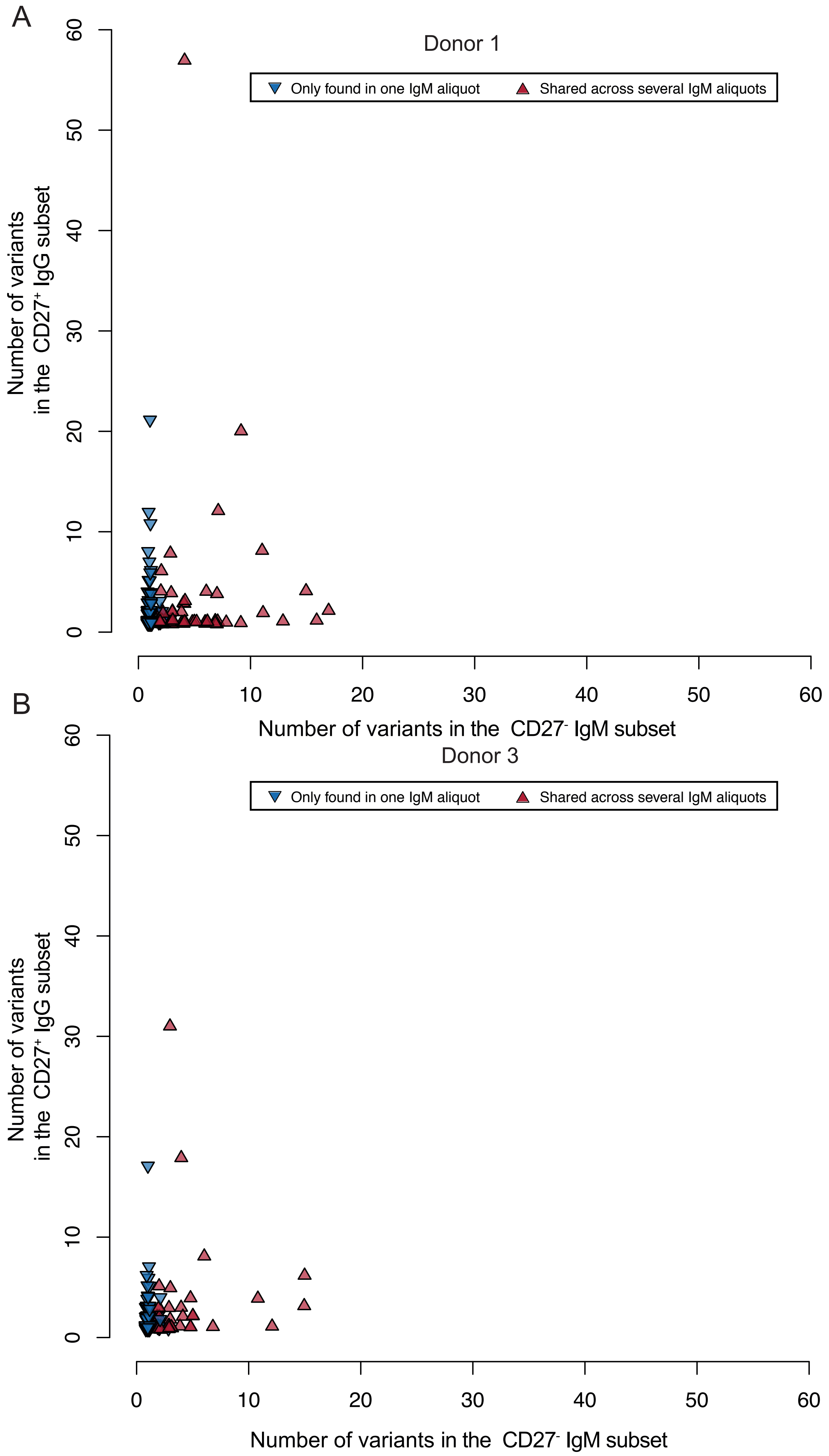


**Figure S4.** Clonal composition of each clonotype that is shared between the IgG and IgM subsets in terms ofits IgG and IgM variants. The red, upward pointing triangle indicates clonotypes that are expaned in the IgM repertoire, whereas the blue triangle highlights clonotypes which could only be found in one IgM aliquot. Panels (**A**) and (**B**) show the results for donors 1 and 3, respectively.
